# Macroevolutionary consequences of twin neck innovations in deep-sea dragonfishes

**DOI:** 10.64898/2026.06.21.733442

**Authors:** Elizabeth Christina Santos, Jonathan Huie, Alessio Capobianco, Rose Faucher, Todd Clardy, William B. Ludt, Giorgio Carnevale, Dahiana Arcila, Christopher M. Martinez

## Abstract

The origin of novel phenotypes can influence access to new ecological resources, which may have positive, neutral, or negative effects on subsequent phenotypic diversification. In this study, we tested the macroevolutionary consequences of a pair of putative functional innovations occurring in deep-sea fishes of the order Stomiiformes. Integrating phylogenetic comparative methods, micro-CT scans, and external body measurements, we recover a mosaic of diversification trends associated with these innovations. We found some evidence for elevated evolutionary rates in tooth morphology associated with the predatory dragonfishes, which possess a gap between their vertebral column and skull that exposes the notochord and enables neck-like flexibility. However, a second novelty building upon the first, a functional neck joint enabling extreme cranial kinesis, was linked to faster rates of skull evolution. Our results suggest that innovations that help shift ecological roles and overcome functional constraints related to those roles, like gape-limitation in prey depauperate habitats, may play an important role in promoting phenotypic diversification. This work builds on a growing body of evidence highlighting how the deep sea promotes phenotypic diversity, generating the extreme forms that are celebrated by scientists and the public alike.

## INTRODUCTION

The evolution of novel traits has long been believed to alter the trajectory of phenotypic diversification. A classic view is that innovations permitting access to new ecological niches, such as powered flight in tetrapods, will increase diversification as the lineage evolves to maximize exploitation of these new resources [1,2]. However, innovations may also have neutral or negative effects on subsequent diversification. For example, if the innovation enhances the lineages’ ability to exploit a single resource, then the performance of the trait will be under strong stabilizing selection. This will promote specialization instead of diversification, constraining phenotypic diversity [3]. As such, the macroevolutionary effects of innovations should be assessed in a hypothesis testing framework rather than assumed [4–7].

The deep sea is thought to impose strong selection pressures on organisms to maximize limited food resources in an environment lacking *in-situ* primary productivity [8,9]. Predatory deep-sea fishes repeatedly evolved characteristics such as large gapes, sharp fangs, and extensible stomachs allowing them to consume large prey if the opportunity arises [10–14]. These observations would suggest that the adaptive landscape of deep-sea environments necessarily constrains phenotypic diversity. For example, recent studies show that jaw lengths are limited to larger sizes in deep-sea fishes compared to shallow-water counterparts [15], but within this constraint the feeding systems of predatory deep-sea fishes still show a broad diversity of shapes and functions [16–19].

Deep-sea fishes display spectacular forms that capture the public’s imagination, surpassing boundaries of what is considered normal across fishes. The focus of this study is the evolutionary consequences of a pair of putative adaptations found in members of the barbeled dragonfish family Stomiidae, specifically (1) a neck-like adaptation characterized by a gap between the neurocranium and first vertebra that exposes the flexible notochord and increases cranial mobility, and (2) a functional joint between the head and anterior trunk region [20]. While all stomiids possess the first innovation [21,22], only a subset of species additionally display the second condition, in which the anterior notochord folds inward to rest beneath the skull and unfolds when the cranium is rotated during gape expansion. This anatomical configuration enables extreme dorsal cranial rotation, helping to increase the overall gape size within the family [20,23,24]. Prior to the comparative anatomy work of Schnell and Johnson [20], the only known fish with a functional neck was *Lepidogalaxias salamandroides,* a freshwater species that uses enhanced cranial mobility to burrow into mud [25] and remains unique among fishes in the ability to turn their head laterally (left and right). More recently, neck-like mobility (i.e., the dynamic contribution of anterior vertebrae to cranial movements) has also been observed in fishes as distantly related as trout (Salmonidae) and frogfish (Antennariidae) [26]. Still, the innovations found in stomiids appear to represent novel evolutionary solutions for a deep-sea context, allowing this lineage to overcome ancestral limitations on cranial mobility and gape size.

In their description of the stomiid neck joint, Schnell and Johnson [20] hypothesized that the adaptation was a phylogenetically informative character, and that two genera with an intermediate condition, *Bathophilus* and *Grammatostomias,* are evidence of stepwise evolution towards full mobility in more derived genera. Innovation may beget innovation as the five genera with the jointed neck showcase other novelties, including transparent teeth [27], the only known case of red bioluminescence in vertebrates [28,29], and the loss of a mandibular membrane or ‘floor’ to the mouth [23,24]. While these special traits qualitatively contribute to the overall diversity of deep-sea fishes, it is less clear whether their evolution has influenced phenotypic diversification of structures across the body plan, contributing to recent results of high diversity of deep-sea fishes compared to shallow-water relatives [15,16,19,30,31].

Here we ask whether the evolution of two neck innovations influences phenotypic diversification across the order Stomiiformes, a group containing the stomiids (barbeled dragonfishes and viperfishes), as well as other common deep-sea clades (e.g., bristlemouths and hatchetfishes). The stomiiform fishes comprise an exclusively deep-pelagic lineage, serving critical ecological roles in midwater communities [13,17,32–34]. Following a recent trend to investigate multiple structures evolving under different selection pressures [35,36], we investigated the associations of the neck innovations with diversification of traits linked to locomotion (body shape) and feeding (overall skull, neurocranium, oral jaws, and teeth). The hypothetical direction of this effect is unclear. On one hand, the neck has been hypothesized to influence diversification in other vertebrates by increasing modularity via segmentation and localized specialization of the body plan [6]. On the other hand, these innovations in stomiids may drive ecological specialization as apex predators in midwater ecosystems and therefore constrain morphology [3,4]. Furthermore, the effect on diversification, if any, may differ by anatomical trait system.

## METHODS

### Phylogeny and divergence time estimation

We based our analysis on a recently published phylogenomic dataset of Stomiiformes [37] comprising 936 nuclear exons (314,607 bp) for 60 stomiiform species (representing 31 of the 52 recognized genera and all four traditionally recognized families) plus four outgroup taxa, with taxon sampling subsequently expanded using mitochondrial CO1 sequences to 135 stomiiform species (of 464 valid species). This is, at present, the best-sampled phylogeny of Stomiiformes with 29% of species sampled. A time-calibrated phylogenetic tree of Stomiiformes was newly estimated in a Bayesian framework using the software RevBayes v.1.3.2 [38], implementing the fossilized birth-death (FBD) model [39,40] (details of this analysis in Supplementary Text 1). The original alignment was reduced to a subset of 37 nuclear exons to improve computational feasibility, totaling 12,420 base pairs, plus CO1 to broaden taxonomic sampling as in the earlier study [37]. These 37 exons were selected based on taxon occupancy, retaining loci with the least missing data while covering the largest number of species, to reduce the effect of missing data on divergence time estimation. In addition, 20 fossil representatives of Stomiiformes were newly included to provide chronostratigraphic data to inform the age of the tree (fossil calibration details in Supplementary Text 2). As the output of FBD analysis, a maximum clade credibility tree was produced to summarize the posterior distribution of trees, with 10% burn-in. Extinct species were pruned from the summary tree before running subsequent phylogenetic comparative analyses.

### Neck innovation groups

Following descriptions from refs [20–22], we delimited three groups for comparative analyses, defined by anatomy posterior to the skull. First, the ancestral condition consists of a stiffened head and anterior trunk region typical of nearly all fishes. Here, the first vertebra attaches directly to the posterior of the neurocranium at the basioccipital. This group includes all stomiiform families exclusive of Stomiidae. The second group contained stomiid genera with a gap between the skull and anterior vertebrae that exposes the notochord, affording additional flexibility to the head and improving cranial rotation during feeding [21,22]. Species in this group, like nearly all stomiids, are midwater predators feeding primarily on other fishes [17,41]. The final group characterized stomiids in which the notochord folds against the posterior of the skull at rest and unfolds during gape expansion, though with some intra-group variation in the degree of notochord folding (see further description below). This improves cranial mobility even further and acts as a functional neck-like joint [20]. Hereafter, we call these categories the (1) ancestral, (2) simple neck, and (3) jointed neck groups (Table S1). We give the caveat that Schnell and Johnson [20] did not examine the extent of cranial rotation in all stomiid genera, though the specific form of a notochord-folding neck joint is not known outside of the genera listed above.

The presence of a postcranial neck-like gap is synonymous with the family Stomiidae, but the evolution of the jointed versus simple neck is potentially more complex. To understand the evolutionary history of neck conditions, we used ancestral state estimation on our expanded phylogeny of Stomiiformes (n=132 species minus outgroups). Here, we simulated 100 potential histories of the evolution of these three states using the ‘make.simmap’ function in phytools v2.5–2 [42,43]. We first compared the fit of equal rates, symmetrical, and all rates different transition models using the ‘fitMk’ function and used the best fitting model in SIMMAP analyses based on a likelihood ratio test.

A secondary question concerns the evolutionary sequence within the jointed neck state itself. Using an earlier phylogeny [29], Schnell and Johnson [20] proposed a stepwise transition from the ancestral condition to an intermediate state with a partially folded notochord, and finally to the fully folded notochord (we include both states in the ‘jointed neck’ term herein). To test this hypothesis, we ran a second ancestral state estimation, as above, with a new grouping scheme: non-joint taxa, partial joint (*Bathophilus* and *Grammatostomias*) and full joint (*Aristostomias, Eustomias, Malacosteus, Pachystomias,* and *Photostomias*).

### Acquiring phenotypic data

Phenotypic data for comparative analyses were measured from fluid-preserved museum specimens of adult stomiiforms and came in three forms: linear measurements of externally visible characteristics of body shape, geometric morphometrics of skulls viewed using micro-CT scans, and linear measurements of teeth viewed using micro-CT scans. Additionally, we extracted subsets of landmarks restricted to the oral jaws and neurocranium, respectively (details below). In summary, downstream comparative analyses were run on five phenotypic datasets representing broad to localized anatomical scales: body plan, whole skull, neurocranium, oral jaws, and teeth.

Taxonomic sampling targeted for phenotypic datasets was designed to match tips in the published phylogeny. Of the 52 genera of Stomiiformes, 42 (80.7%) were represented in both the phylogeny and phenotypic datasets, including all seven genera possessing a jointed neck. Of the remaining 10 stomiiform genera, two were sampled in the tree but missing from body shape and/or CT scan datasets (*Rhadinesthes*, *Valenciennellus*), two had CT scans but not sampled in the tree (*Pollichthys, Trigonolampa*), and six were not found in any datasets (*Araiophos, Chirostomias, Eupogonesthes, Manducus, Sonoda, Thorophos*).

#### Body shape

External body shape was measured using handheld digital calipers with a minimum resolution of 0.1 mm. We took eight measurements following [44]: standard length, maximum body depth, maximum fish width, head depth, lower jaw length, mouth width, minimum caudal peduncle depth and minimum caudal peduncle width. Five additional measurements were taken, including: maximum eye diameter, perpendicular eye diameter, minimum interorbital distance, head length, and caudal peduncle length. Raw body shape measurements and associated specimen catalog numbers can be found in the Dryad data repository. Body shape was measured from 260 specimens representing 73 species (1–5 fish measured per species based on specimen availability and condition; mean 3.56 specimens) and 43 stomiiform genera.

Measurements were scaled using log-shape ratios following [15,19,44]. Each variable was divided by the geometric mean of standard length, maximum body depth and maximum fish width, and then log-transformed. We then took the mean of all individuals for a given variable to represent the species in our analyses. Variables were centered around a mean of zero and scaled to unit variance using the ‘scale’ function in base R, then exported for some downstream analyses. To visualize body shape diversity, principal component analysis (PCA) was run using the ‘prcomp’ function in geomorph.

#### Skeletal anatomy

Skull shape was measured using three-dimensional geometric morphometrics based on micro-CT scans of skulls. Our CT scan dataset spanned 44 genera and 78 species (n=1 scan per species). Of the 78 scans, 69 were newly generated for this study using a Bruker SkyScan 1273 at the Natural History Museum of Los Angeles County. Voucher museum specimens for these new scans were from the following institutions (museum acronyms from [45]): Harvard Museum of Comparative Zoology (MCZ), Natural History Museum of Los Angeles County (LACM), Scripps Institution of Oceanography (SIO), and the University of Florida (UF). An additional nine scans were downloaded from MorphoSource [46,47]. See table S1 for all scan metadata.

Skulls were segmented and digitized with 102 landmarks (42 fixed and 60 sliding semi-landmarks) using 3D Slicer v. 5.6.1 [48]. Our landmark scheme was modified from ref. [19] to adapt it to anatomy of non-acanthomorph fishes (Table S2, Fig. S1). Landmarks were treated as bilaterally symmetrical and only placed on the left side of the skull. To reduce the influence of preservation artefacts and postural variation in skull shape, we performed a local superimposition [49]. This approach allowed us to standardize the position of eight individual skull elements: premaxilla, maxilla, anguloarticular, dentary, neurocranium, hyomandibula, hyoid complex, and palatine. For two additional sets of analyses, we extracted subsets of landmarks placed on the neurocranium (18 landmarks) and oral jaws (43 landmarks), respectively (Fig. S1). In these cases, we aligned landmarks using generalized Procrustes analysis of the neurocranium, and localized Procrustes superimposition analysis including the four bones composing the oral jaws (premaxilla, maxilla, anguloarticular, and dentary), respectively. The function ‘gpagen’ from the geomorph v. 4.0.10 R package [50,51] was used for shape alignments, paired with custom R scripts for local superimposition [19,49]. To visualize phenotypic diversity, we ran a PCA using the ‘gm.prcomp’ function in geomorph, inputting each set of aligned coordinates (whole skull, neurocranium and oral jaws).

We also used micro-CT scans to make linear measurements of teeth in 3D Slicer. The following measurements were taken from the lower jaw: length of the longest tooth (from tip to the base), width of the longest tooth taken at the base, aspect ratio of the longest tooth calculated as tooth length divided by tooth width, in-lever length of the primary jaw closing muscle (*adductor mandibulae*) measured as the distance from the quadratomandibular joint to the distal tip of the coronoid process of the articular, length of the lower jaw from the quadratomandibular joint to the anterior tip of the dentary, and jaw closing mechanical advantage calculated as in-lever divided by jaw length (Fig. S2). Tooth length and width were scaled by dividing by lower jaw length. Four morphological traits were constructed from the above measurements: size-corrected tooth length and width, tooth aspect ratio, and jaw closing mechanical advantage (Fig. S2). Traits were centered and scaled using the ‘scale’ function in R, then exported for downstream analyses. To visualize tooth shape diversity, we also ran a PCA using the ‘prcomp’ function in geomorph.

### Evolutionary rate and disparity analyses

Not all comparative analyses were phylogenetic in nature. PCA was run on each data subset prior to pruning taxa not found in the phylogeny, as this allowed us to include 12 additional species with CT scans. Likewise, landmarks were aligned using the full 78-species dataset (Table S1). For all phylogenetic comparative analyses, we pruned the CT scan-based datasets for congruence with the phylogeny (n=66 species). The body shape dataset did not require pruning because all 73 species with measurements were also sampled in the phylogeny.

Rates of multivariate trait evolution were inferred using two approaches. First, we used the ‘compare.evol.rates’ function in geomorph to compare Brownian motion rates for the three groups (ancestral, simple neck, and jointed neck). We ran analyses separately for the five phenotypic datasets (body shape, whole skull, neurocranium, oral jaws, and teeth). We also compared phenotypic disparity (i.e., multivariate variance) across the three groups using the ‘morphol.disparity’ function in geomorph. We prefer to interpret disparities alongside evolutionary rates, as the latter provides important temporal context when interpreting standing phenotypic diversity [52,53]. While the morphological disparity analysis is non-phylogenetic, we used the pruned datasets to ease comparison with rates. For both functions, the inputs were scaled variables or aligned shape coordinates, and significance was based on 10,000 simulations (rates) or permutations (disparity).

Second, we tested for shifts in evolutionary rates on the phylogeny using a phylogenetic ridge regression implemented in the RRphylo package v.3.0.2 [54]. Unlike ‘compare.evol.rates’, this approach does not require an *a priori* hypothesis of the location of shifts nor user-defined groups to be compared. We first used the ‘RRphylo’ function to perform phylogenetic ridge regression, inputting PC axes to represent phenotypic variation, following ref. [55]. Next, we used the function ‘search.shift’ in auto-recognize mode to detect whether each clade in the tree represented a significant rate shift (either increasing or decreasing) relative to background diversification across the tree.

## RESULTS

### Divergence time estimation

Our divergence time analysis using the FBD approach recovers crown Stomiiformes as originating late in the Early Cretaceous, around 116.6 Ma (95% highest posterior density (HPD): 137.2–91.4 Ma). Dragonfishes (Stomiidae) appear towards the end of the Late Cretaceous, around 76.9 Ma (95% HPD: 93.5–60.3 Ma), while the dragonfish subclade characterized by the evolutionary innovation of the neck joint (comprising *Aristostomias*, *Bathophilus*, *Eustomias*, *Grammatostomias*, *Malacosteus*, *Pachystomias*, and *Photostomias*, plus the non-jointed genus *Idiacanthus*) originated in the early Eocene, around 52.7 Ma (95% HPD: 65.4–40.2 Ma). Most extant stomiid genera seem to have been already established before the end of the Oligocene (23 Ma). FBD tree with fossil placement and posterior densities is shown in Fig. S3.

### Evolution of the jointed neck innovation

The best-fitting transition model was equal rates for the neck states, and symmetrical rates for the degree of notochord folding within the jointed neck (Table S3). Estimates of ancestral neck states showed two origins of the simple neck condition, one at the base of the family Stomiidae and a second in the genus *Idiacanthus*, the latter a reversal from the jointed neck state (Fig. S4). Additionally, a single origin of the jointed neck condition was recovered within the stomiids. A separate analysis to examine the sequence of intermediate (partially folded) and fully jointed (fully folded) neck states showed that, contrary to previous hypotheses [20], the innovation did not evolve in a stepwise fashion. Rather, the partially folded state evolved twice, independently in *Bathophilus* and *Grammatostomias*, from lineages with the fully folded state (Fig. S5).

### Shape diversification across neck types

Morphospaces for anatomical regions considered in this study are shown in Fig. 2. Rates estimated using ‘compare.evol.rates’ are plotted against disparity in Fig. 3; absolute values of both are given in Table S4. Significance of rates and disparity are given in Tables S4 and S5.

**Figure 1.**
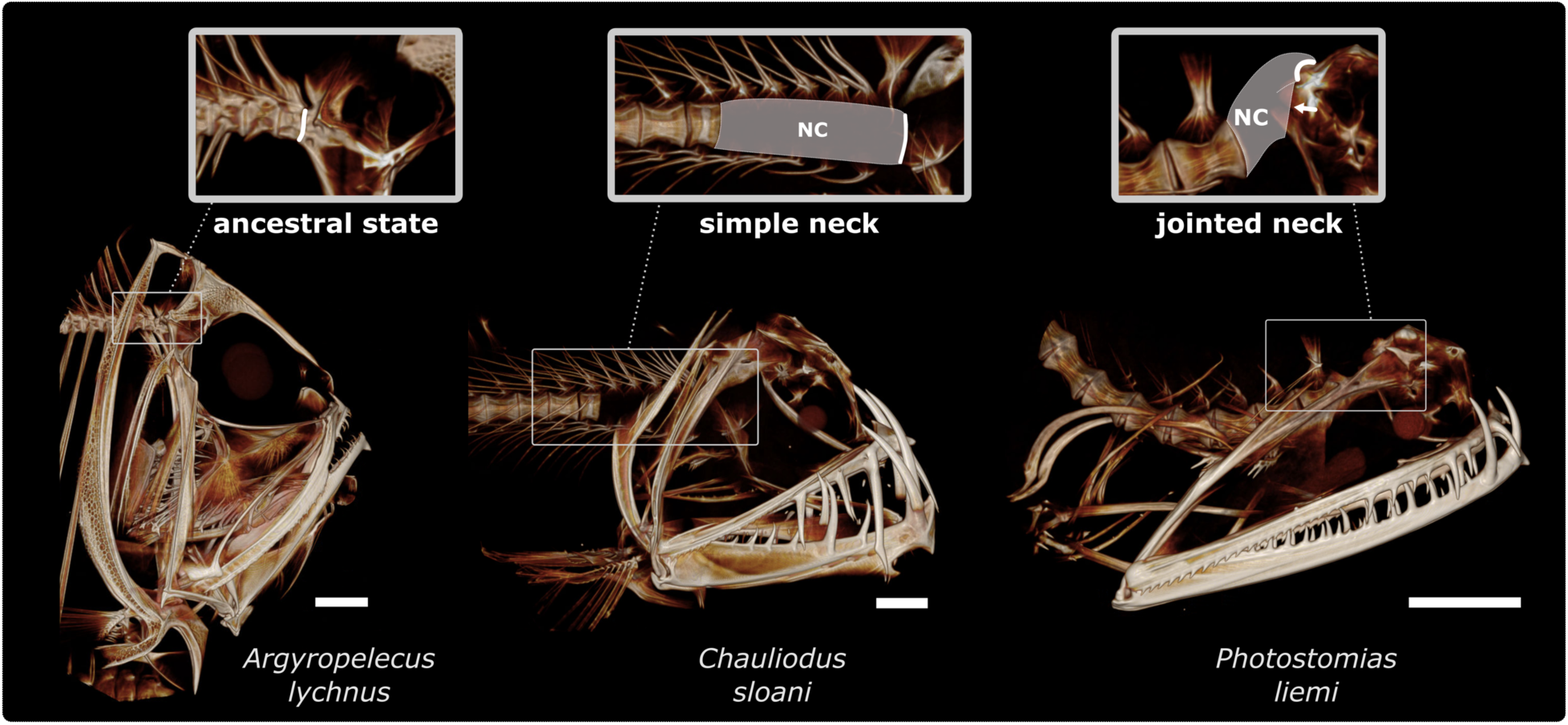
Representative taxa from three functional groups examined in this paper. Micro-CT scans are shown to visualize skeletal morphology, with white scale bars = 5mm. Insets show the anterior vertebrae with superimposed drawings of exposed notochord (NC) and the site of connection to the neurocranium (bold white line) based on ref. [20]. The ancestral, neckless, condition in Stomiiformes (left) is common to nearly all other fishes, consisting of the anterior vertebral disc articulating directly with the back of the skull. Species with a flexible notochord-supported gap between the skull and vertebrae form the “simple neck” group (middle), and those with the gap plus folding of the notochord against the base of the skull make up the “jointed neck” group (right).

**Figure 2.**
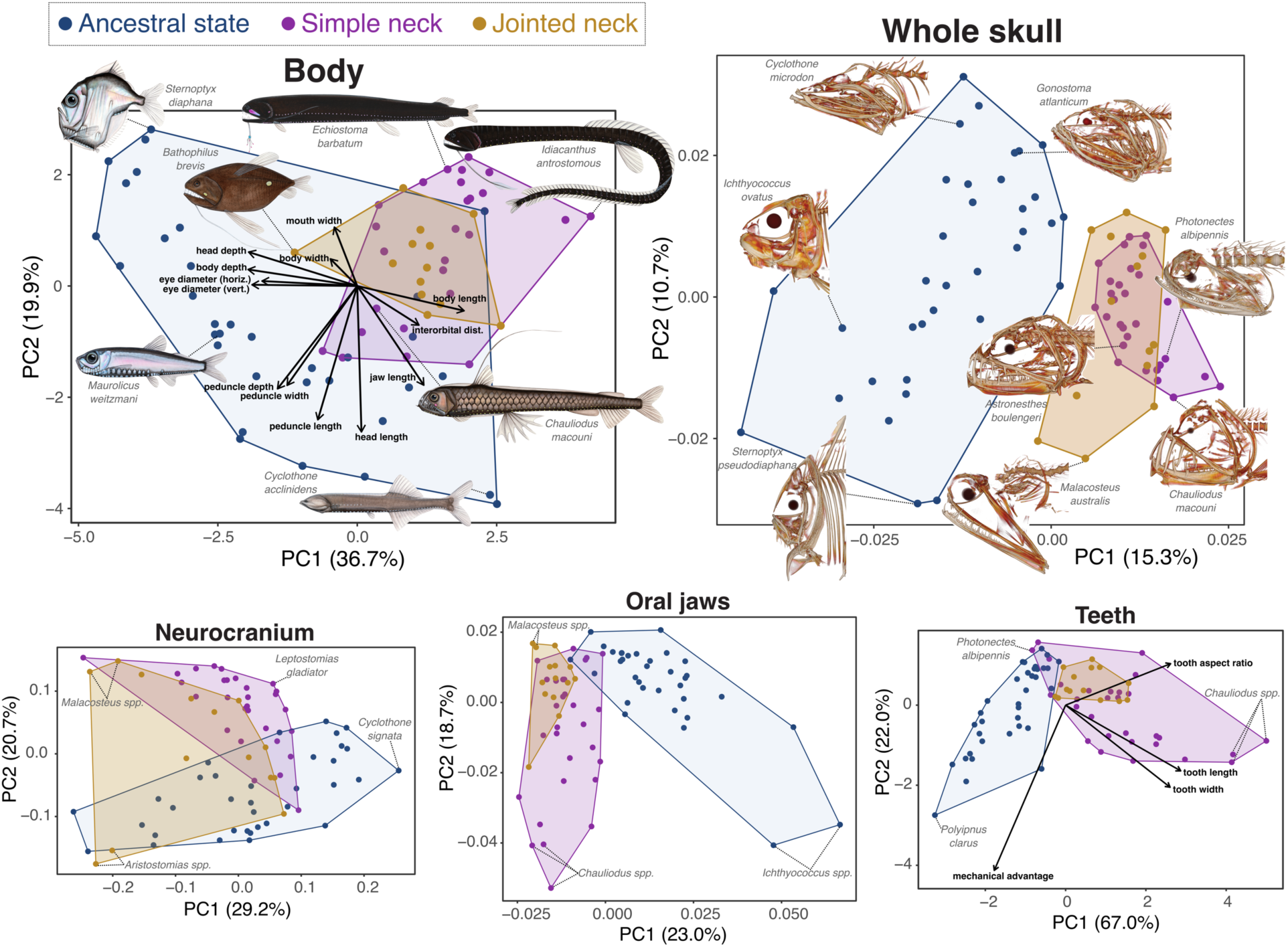
Morphospaces of Stomiiformes by phenotypic subset, represented by PCs 1 and 2 from PCAs. Shapes of the skull, neurocranium, and oral jaws are based on geometric morphometrics, while body and teeth shape are based on linear trait measurements. Images of overall body shape and skull morphology (based on micro-CT scans) illustrate phenotypic variation across major axes of diversity.

**Figure 3.**
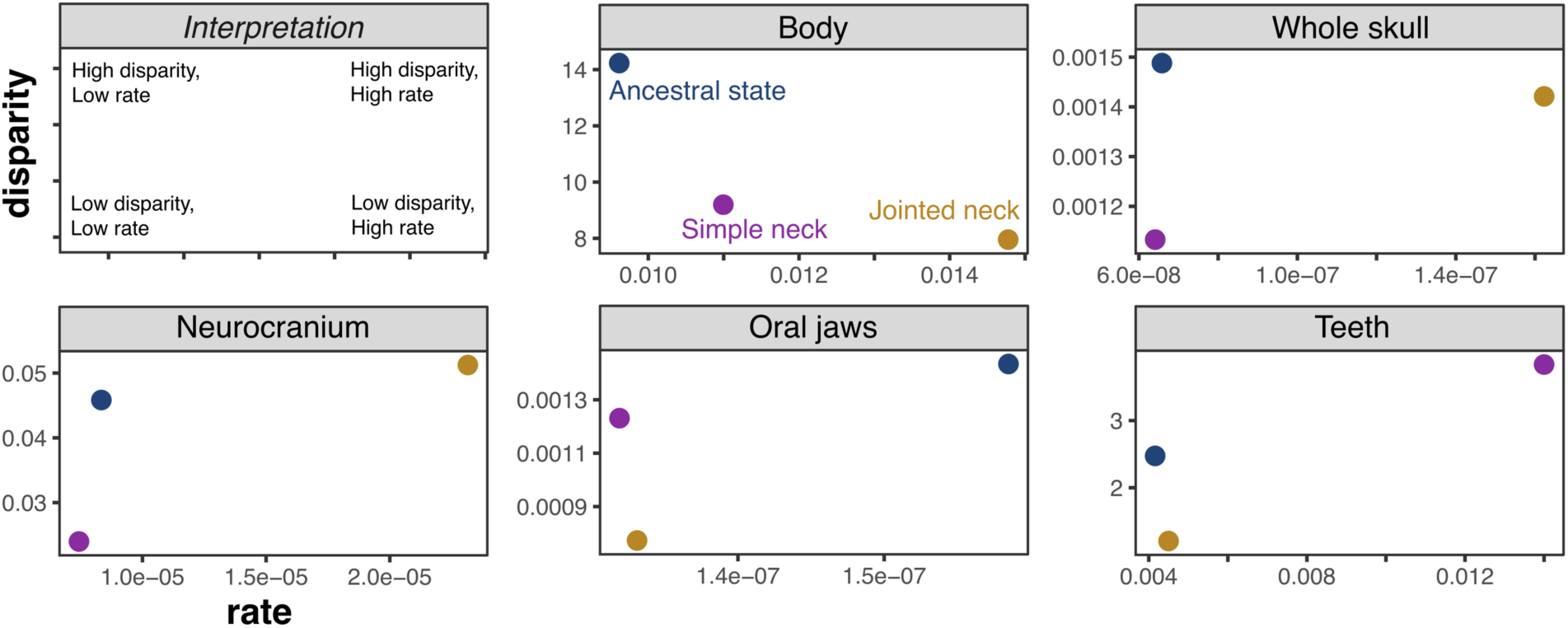
Multivariate Brownian motion rates of shape evolution plotted against morphological disparity estimates for each functional group. The upper left panel assists with interpretation. Values of rates and disparity are given in Table S4.

Rate shifts estimated using RRphylo are shown in Fig. 4. Below we discuss results for each of the five phenotypes individually, in order of anatomically broad to localized.

**Figure 4.**
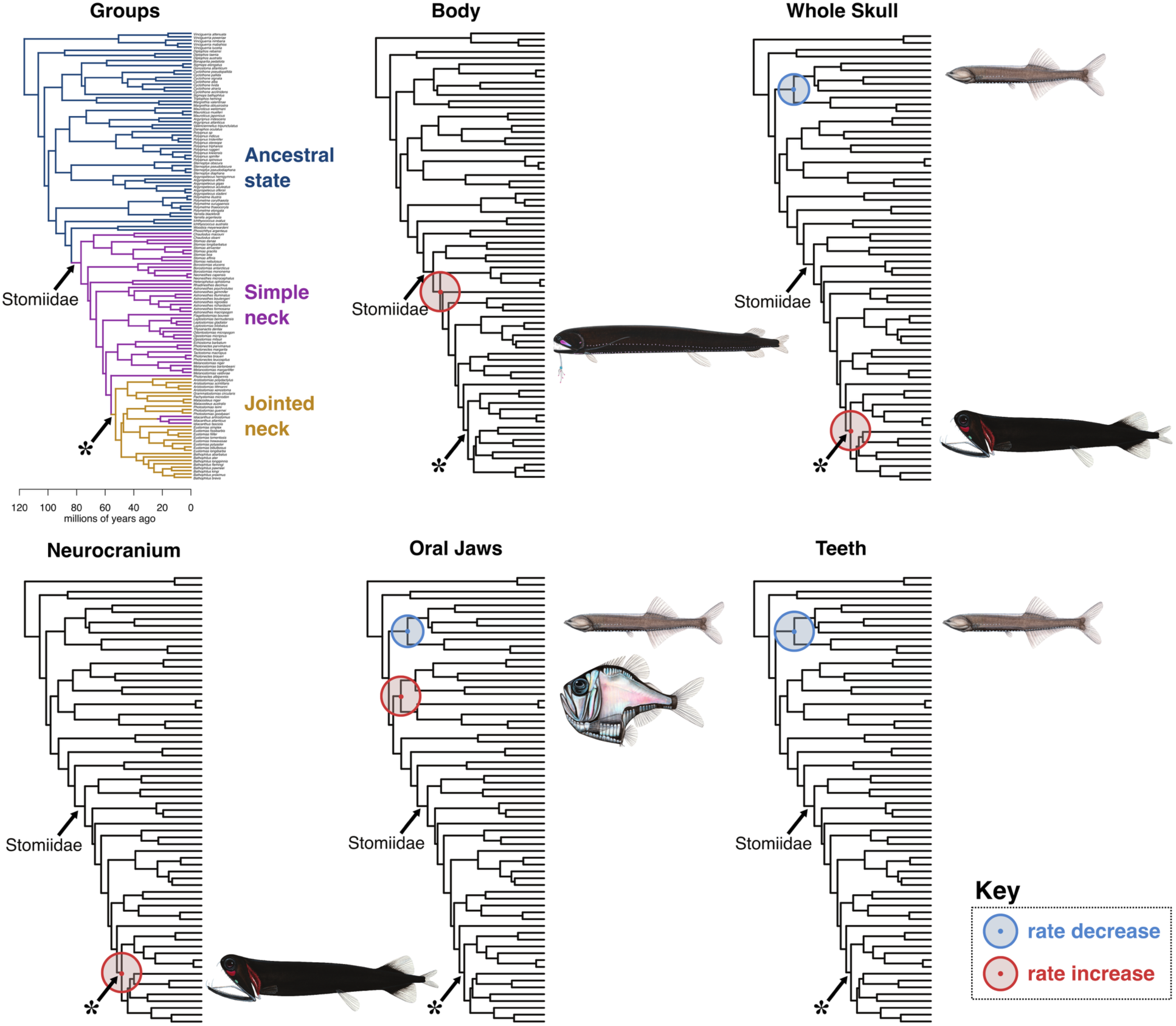
Shifts in the rate of shape evolution estimated using group-blind analyses in RRphylo. Nodes with a significant rate shift are indicated with a blue (rate decrease) or red (rate increase) dot. Within each dataset, the size of the circle surrounding the node is scaled by the magnitude of the rate shift. Asterisk indicates the node associated with evolution of the jointed neck based on SIMMAP analyses (Fig. S4). Representative taxa associated with rate shifts are illustrated.

#### Body shape

Stomiiform fishes with the ancestral vertebral type show much wider body shape diversity than the stomiids, as illustrated by the extremely laterally compressed hatchetfishes, stomiid-like lightfishes, and wispy bristlemouths. This functional group has higher disparity, but lower rates of body shape evolution compared to both stomiid groups with neck innovations (Fig. 3). Body shape diversity of stomiids with jointed necks is nested within the simple necked stomiids in morphospace, except for *Bathophilus brevis* which has a truncated body plan unique within Stomiidae and even within its own genus (Fig. 2). Evolutionary rate shift analyses show a rate increase at a node within Stomiidae subtending most of the genera in the family but not associated with any known trait shared by these taxa (Fig. 4).

#### Whole skull

The primary axis of skull shape evolution reflects changes in overall cranial aspect ratio, from a laterally compressed shape on the negative extreme of PC1 associated with hatchetfishes to a more streamlined shape with long jaws on the positive extreme, associated with dragonfishes of both simple and jointed neck functional groups (Fig. 2). The jointed neck group forms a larger, though partially overlapping, distribution in skull-shape morphospace relative to the older simple neck group. There are no significant differences in disparity, and the rate of skull shape evolution in the jointed neck group was over 2-fold greater than the other two groups (Fig. 3; Tables S4, S5). This result is supported by group-blind rate shift analyses, which show a significant increase in rates at the node corresponding with the single evolutionary origin of the folded notochord, as inferred by SIMMAP (Fig. 4). Additionally, these same analyses also show a rate decrease in skull shape associated with Gonostomatidae (bristlemouths).

#### Neurocranium

The neurocranium is the only phenotype examined for which the jointed neck group has both the highest standing disparity and highest evolutionary rate (Tables S4, S5). Here, diversification of species with a jointed neck was 3.1-fold greater than the simple neck group and 2.8-fold higher than those with the ancestral condition. Rate shift analyses corroborate these results, showing a rate increase at the node associated with the origin of the joint. Changes in the neurocranium along PC1 can be described as a shift from foreshortened and deep neurocrania with large orbits (hatchetfishes) on the negative end, to shallow and elongate with small orbits (bristlemouths) on the positive end (Fig. 2). Stomiids are widespread in the morphospace, from intermediate to low values of PC1. Within this family, the jointed neck group occupies both extreme positions along PC2, associated on the negative end with a terminally directed supraethmoid relative to the orbits (*Aristostomias*), and on the positive end with a domed, forward-facing orbit and ventrally deflected supraethmoid (*Malacosteus*).

#### Oral jaws

The primary axis of jaw shape morphospace is associated with elongate jaws on the negative end (dragonfishes) and short but robust jaws (lightfishes) on the positive end (Fig. 2).

Stomiids were limited to negative values of PC1 but displayed a wide range of shapes along PC2, which was associated with gape angle and size of the lower jaw relative to upper jaw. Jaws in the jointed neck group occupy a limited area of morphospace associated with positive PC2 values, mostly nested within simple-necked dragonfishes, with the exceptionally elongate and anteriorly-directed jaws of *Malacosteus* and *Pachystomias* representing extreme forms. There were no significant differences in disparity or evolutionary rates detected. Jaw shape evolved at similar rates in stomiids with simple and jointed necks, though the former shows higher standing disparity, presumably reflecting its older age and greater time to diversify (Fig. 3). RRphylo detected two rate shifts in jaw shape (Fig. 4), both outside of Stomiidae: a rate decrease associated with Gonostomatidae (bristlemouths) and an increase associated with Sternoptychidae (marine hatchetfishes).

#### Teeth

As with the oral jaws, teeth of jointed neck taxa form a constrained subset of the morphospace occupied by simple necked stomiids (Fig. 2). This corresponded with the simple neck group containing 3.2 times greater disparity of tooth traits, and 3.1 times faster diversification than the jointed neck group (Fig. 3; Table S4). However, this same result was not recovered by the group-blind rate shift analysis. Mechanical advantage and tooth aspect ratio load in nearly opposite directions, with neckless taxa possessing higher mechanical advantage values. Only a single rate shift was detected (Fig. 4): a reduced rate of tooth evolution in Gonostomatidae (bristlemouths).

#### Summary

The jointed neck innovation is associated with an increase in evolutionary rates and morphological disparity in skull shape, especially the neurocranium. In contrast, we found no evidence that the jointed neck is associated with elevated rates of jaw or tooth evolution relative to stomiids with a simple neck. Rather, taxa with the jointed neck appear to be largely constrained with respect to these traits. Outside of the stomiids, we found shifts towards lower rates of skull, jaw and tooth shape evolution in the bristlemouths and a shift towards faster jaw shape evolution in the hatchetfishes. Finally, body shape evolution shows a unique pattern where the ancestrally neckless taxa have the highest disparity but the lowest evolutionary rates.

## DISCUSSION

In this study, we assessed the macroevolutionary consequences of two modifications of the anterior vertebral column in deep-sea fishes from the order Stomiiformes. Each modification enabled a degree of neck-like mobility of the cranium, augmenting a critical function for fish feeding [26]. We approached this system with the understanding that evolutionary innovations can have varied effects on the diversification of other phenotypes based on their degree of anatomical interaction with the novel trait and the prevailing mode of selection [3]. A major finding of this study was that elevated phenotypic diversification of the skull, and particularly the neurocranium, was most prominent in stomiids containing both neck innovations (a flexible postcranial vertebral gap, and a folded notochord acting as a cranial joint), but not for those with just a postcranial gap (“simple neck” herein). This result mirrors patterns observed in parrotfishes, where increased rates of morphological evolution were recovered in species possessing a novel lower jaw joint plus a modified pharyngeal jaw, but not taxa with the pharyngeal adaptation alone [56], highlighting the synergistic effects of sequential innovations on diversification. Beyond the neurocranium, our results pointed to a patchwork of patterns of evolution for different trait systems related to feeding and locomotion in an ecologically prominent lineage of deep-sea fishes.

### Macroevolutionary effects of twin neck innovations

A flexible notochord-supported gap between the skull and first vertebral element in stomiid fishes led to improved cranial rotation by acting as a simple neck. Since all stomiids possess this general functional trait [20–22], the evolution of the simple neck likely coincided (along with bioluminescent chin barbels) with the origin of a clade consisting almost exclusively of apex midwater predators [13,17,32]. This is consistent with the comparably high rates of tooth evolution in the simple neck group (Fig. 3; though not found in our group-blind rate shift analyses), which contains species with the largest fangs encountered among stomiiform fishes (Fig. 2). We should reiterate that we use the term “simple” primarily to distinguish from taxa that additionally possess a folded notochord. In reality, the simple neck is expressed by a variety of anatomical manifestations, including unique modifications in taxa like *Chauliodus* and *Stomias*, the latter even described as possessing a “pseudo-craniovertebral articulation” [21,22].

Within stomiids, the evolution of a neck joint leveraged the previous innovation of a gap between the vertebral column and cranium to further enhance rotational freedom at a discrete hinge at the base of the skull. The result is the greatest achievement of cranial elevation (up to ∼80° of rotation) ever observed for bony fishes [20]. Along with this second innovation came higher diversification rates of the neurocranium, which seems fitting as it contains the connection to the flexible notochord and joint site enabling extreme mobility. The lability of neurocranial morphology in this group is exemplified by the large spread in morphospace between related ‘loosejaw’ taxa (Fig. 2). The neurocranium of *Malacosteus* is especially unique for its domed “forehead” with large forward-facing orbits, such that past authors suggested that this genus has binocular vision [24]. Other authors mused that the neurocranium of *Malacosteus* does not look like it belongs to a fish [57]. However, not all cranial features in these species showed elevated rates. Rather, jaws were highly constrained around long, gracile forms with weak mechanical advantage and moderate tooth size (Fig. 2 D&E). Such patterns could reflect stabilizing selection for large gapes and high cranial mobility (a trade-off with strong bite force). These results are consistent with previous literature suggesting that the loosejaw condition of some jointed neck taxa (i.e., the loss of a mandibular membrane; Table S6) reduces drag on the jaws, circumventing a trade-off and permitting evolution of a larger gape than other stomiids [23]. The collection of extreme adaptations found in the jointed neck stomiids appear to have evolved in the context of overcoming gape limitation in low-food conditions, which is a major constraint for predatory fishes [58,59].

An innovation is predicted to promote phenotypic diversification if it permits ecological diversification through niche segmentation [3]. While lifestyles of deep-sea fishes are difficult to assess, one study documented four dietary guilds among stomiids based on gut contents [32].

Two of the four guilds were restricted to taxa with the jointed neck innovation, suggesting that this group expanded the dietary repertoire of the family. These taxa may also show behavioral differences from other stomiids including reduced vertical migration and active pursuit of prey illuminated by red bioluminescence [24,60]. Changes in the neurocranium associated with orbit size and position of the ethmoid region, even within the jointed neck group, may indicate differences in gape position and sensory modalities, further hinting at ecological diversification.

### Evolutionary dynamics among ancestrally neckless taxa

Phenotypic rate shifts were not exclusive to stomiiforms with neck-like innovations. Taxa with the ancestral, neckless form of anterior vertebrae are highly diverse in their own right, also displaying a range of extreme morphologies (Fig. 2). The marine hatchetfishes (Sternoptychidae) show elevated rates of oral jaw diversification (Fig. 4), presumably related to massive cranial reorganization during the acquisition of their uniquely deep-bodied and laterally compressed body plans.

Among stomiiforms, bristlemouths (Gonostomatidae) buck the general trend with consistent rate shifts towards slower diversification for the skull, lower jaws, and teeth. These shifts were all located at same node at the base of the family (Fig. 4) and are aligned with observations that bristlemouths have a strong affinity towards planktonic crustaceans, accounting for as much as 91% of their dietary biomass [41]. Fossil bristlemouths show evidence of having a similar diet to modern species[61] and microfossils document long-term conservatism of tooth morphology in this family [62]. Despite the overall conservatism of their skulls, unique cranial novelties have been identified, including a spiral-patterning on the teeth of *Cyclothone* that is hypothesized to snag like Velcro on the antennae of small crustacean prey [62].

### Implications for systematics

We found that the folded notochord neck joint evolved once, with two transitions to an intermediate state and one back to the simple neck state (Figs. S4, S5), contradicting previous hypothesis of stepwise evolution [20]. The origin of the jointed neck coincides with a clade that is rife with innovations, including the loosejaw condition and red bioluminescence [23,29], that are found in no other vertebrates (Table S6). This suggests that the folded notochord may have set the stage for additional novelties via runaway selection towards extreme traits for predatory lifestyles. Traditionally, the subfamily Malacosteinae (‘loosejaws’) consists of *Aristostomias*, *Photostomias*, and *Malacosteus*, and this rank name is still frequently used [17,63]. Though molecular phylogenies have never supported the three loosejaw genera as a clade, the seven genera sharing the jointed neck consistently form a clade in molecular [29,37] and morphological [57] phylogenetic analyses. We advocate for the consideration of the jointed neck clade as a subfamily in future taxonomic studies, which would address the non-monophyly of Malacosteinae while recognizing the many innovations endemic to this lineage. Further clarity on the placement of *Idiacanthus*, a simple-necked genus nested in this clade, will be important for resolving the taxonomy of the group.

### Diversification in the deep sea

Patterns of stomiiform evolution provide generalizable insights for deep-sea fish diversification, contributing to a growing body of literature on the topic [15,16,18,19,30,31,53,64]. First, evolutionary innovation continues long after deep-sea colonization. The classic model of adaptive radiation is an early burst of diversification followed by canalization as ecological niches are filled [1,2]. According to fossil and molecular evidence, Stomiiformes have inhabited the deep sea since at least the Early Cretaceous [64,65] (Fig. S3).

Yet, the jointed neck innovation and other subsequent feeding adaptions (Table S6) evolved within a nested lineage that is Eocene in age according to our FBD analyses (Fig. S3). In parallel with stomiids, some anglerfishes evolved an upper jaw novelty as recently as 10 million years ago, despite the lineage as a whole occupying midwater habitats for 60 million years [16,19]. This suggests that ecological saturation has not prevented the origin of novelties in the deep sea, which can seemingly arise at any time. Late-stage innovation may be an underappreciated aspect of evolutionary radiations in general, defying the canonical early burst model and requiring new explanations [36].

Second, a picture is now beginning to emerge in which morphological and functional diversity outpaces ecological variation in the deep-sea [16,17]. Stomiiforms collectively boast impressive diversity across various anatomical scales (Fig. 2). Previous work has identified a trend of increasing body shape diversity at greater ocean depths, such that some deep-sea fishes with similar ecologies and habitats occupy opposite extremes of the same phenotypic axes [15,19]. This complicates understanding of causal drivers of phenotypic diversification. The traditional adaptive radiation model invokes ecomorphological coupling in order to maximize resource exploitation [1,4,35]. Dietary studies of stomiids and other deep-sea fishes show higher resource partitioning than once believed for this food-limited environment, but not enough to fully account for cross-species or cross-genus morphological differences [17,32]. Therefore, it appears that processes and mechanisms used to explain the diversity of shallow-water fishes are likely insufficient to describe patterns observed in deep-sea fishes [15,19,53].

Feeding adaptations abound in the deep sea, where meals can be few and far between. Our work suggests that the evolution of a functional neck joint corresponded with elevated rates of evolution of the neurocranium in certain dragonfishes, presumably linked to an improved ability to exploit scarce dietary resources. In contrast, bristlemouths consistently showed limited diversification across trait systems. Together, these scenarios highlight the complex selection pressures acting on midwater organisms [66]. This research motivates future work on the biomechanical links between different trait systems during integrated feeding behaviors and their underlying patterns of evolutionary integration.

## ACKNOWLEDGEMENTS

We thank Elias Zamora and Nadia Pourshahmir for assistance with digitizing anatomical landmarks on skulls. We are also grateful to Julie Johnson (Life Science Studios) for providing illustrations used to visualize body plan diversity across stomiiforms. Finally, we thank Tina Wu for helping to produce micro-CT scans at LACM. This work was supported by NSF grants DEB-2401223 and DEB-2519905 awarded to D.A. and start-up funds from UCI to C.M.M.

## Notes

### Competing Interest Statement

The authors have declared no competing interest.

