## Supplemental information for "Macroevolutionary consequences of twin neck innovations in deep-sea dragonfishes"

Supplementary Information

Table of Contents:


- Supplementary Text 1: Additional Methods
- Supplementary Text 2: Fossil calibration list
- Figure S1: Cranial landmark placement
- Figure S2: Tooth measurements
- Figure S3: FBD maximum clade credibility tree with extinct taxa
- Figure S4: Ancestral state estimation for neck innovation state using SIMMAP
- Figure S5: Ancestral state estimation for notochord folding state using SIMMAP
- Table S1: Metadata for CT scans and neck innovation category assignments
- Table S2: Description of landmark placements
- Table S3: Likelihood ratio test for transition rate Mk models
- Table S4: Significance of rate differences using compare.evol.rates
- Table S5: Significance of variance differences using morphol.disparity
- Table S6: Summary of additional evolutionary novelties co-occurring with the folded notochord

**Supplementary Text 1:** Additional Methods

*Fossilized birth-death analysis:*

We used a previously published phylogram of Stomiiformes for this study [1]. This phylogeny was originally constructed with 936 nuclear exons from 64 species, plus CO1 sequences to expand taxonomic sampling to 136 extant species in the final phylogram (132 stomiiforms plus four outgroups). This is, at present, the best-sampled phylogeny of Stomiiformes with 28.4% of species sampled. A time-calibrated phylogenetic tree of Stomiiformes was estimated in a Bayesian framework using the software RevBayes v.1.3.2 [2]. The alignment for this analysis was reduced to a subset of 37 nuclear exons for the 64 species with genomic data in order to improve computational feasibility, totaling 12,420 base pairs, plus CO1 for 135 species to broaden taxonomic sampling as in the earlier study [1]. In addition to the 136 extant species, 20 fossil representatives of Stomiiformes were included to provide chronostratigraphic data to inform the age of the tree (fossil details in Supplementary Text 2). The molecular data was partitioned into four subsets: three subsets for the concatenated exon data (one for each codon position), and the fourth subset for the CO1 alignment. Each data subset was assigned its own GTR substitution model, with among-site rate variation modeled by a gamma distribution discretized in 8 rate categories.

The fossilized birth-death process (FBD) was picked as the lineage evolution model [3,4]. We applied an unresolved FBD approach, where there is no morphological character data and the phylogenetic position of extinct species is constrained a priori by topological constraints that can be more or less strict, depending on the confidence in the taxonomic placement of a fossil [4]. Estimated speciation, extinction, and fossil sampling rates were kept constant through time, and their priors were set as uniform distributions bounded between 0 and 10 events per lineage per million years. Sampling fossils as direct ancestors of other sampled lineages (i.e., sampled ancestors) was allowed. Sampling probability of extant species was fixed to 132/464, corresponding to the fraction of extant stomiiform species included in the analysis, and assuming uniform sampling of extant species. A topological backbone was used to fix the relative position of the 64 species with exon data to the genomic topology of Chang et al. [1], while the position of the species represented only by mitochondrial data was newly estimated in the analysis. For this reason, some topological differences between this study and Chang et al. [1] are possible. Constraints were defined to place fossils in the phylogeny based on their taxonomy and systematics (calibration details in Supplementary Text 1). Tip ages of fossil taxa were given a uniform prior distribution ranging from the minimum to the maximum age of the deposit in which each fossil has been found. The prior on the root age of the tree was set as a uniform distribution, with minimum = 94 Ma, corresponding to the age of the oldest fossil included in our analysis, and maximum = 147.6 Ma, corresponding to the maximum of the 95% credible interval of the MRCA of Stomiiformes and Osmeriformes in the phylogenetic analysis of Hughes et al. [5].

As a clock model, we used a relaxed uncorrelated lognormal clock to allow for rate variation across branches. The prior on the mean clock rate was set as a log-uniform distribution ranging from 10-5 to 10-1 substitutions per site per million years, and the prior on the standard deviation of the clock rate was set as an exponential distribution with mean=0.587405 (equivalent to one order of magnitude of clock rate variation across branches).

The Markov chain Monte Carlo was set up as two independent runs, running for 50,000 iterations and sampling every 10, with 1100.2 moves per iteration. Convergence between runs was checked by visually inspecting and calculating effective sample sizes of parameter estimates on Tracer v.1.7.2 [6]. A maximum clade credibility tree was produced to summarize the posterior distribution of trees, with 10% burn-in. Extinct species were pruned from the summary tree before running subsequent phylogenetic comparative analyses. The phylogenetic placement of extinct taxa can be seen in Fig. S3.

**Supplementary Text 2:** List of 20 fossil Stomiiformes used for the divergence time estimation under the fossilized birth-death (FBD) approach

### 1. *Paravinciguerria praecursor*

Constrained phylogenetic position: Stem Stomiiformes (*Vinciguerria nimbaria* + *Photostomias guernei*)

Fossil: *Paravinciguerria praecursor*, MNHN-F DTS 74. This species, known from the Upper Cretaceous organic-rich shales of Jbel Tselfat, Morocco, Cinto Euganeo, Italy, and northeastern Sicily (Floresta and Malvagna), is used to provide a minimum age for the total group Stomiiformes [7].

Age: The black shales of Jbel Tselfat, Cinto Euganeo, Floresta and Malvagna represent the sedimentary expression of the Oceanic Anoxic Event 2 (OAE 2) that occurred around the Cenomanian-Turonian boundary, between circa 94.9 and 93.9 Ma (see, e.g., [8]).

### 2. *Vinciguerria distincta*

Constrained phylogenetic position: Stem *Vinciguerria* (*Vinciguerria nimbaria* + *Vinciguerria poweriae*)

Fossil: *Vinciguerria distincta*, PIN 1413/61. Danil'chenko [9] described this species based on about ten articulated skeletal remains from the laminated organic-rich shales of the Dabakhan Formation exposed along the Dabakhanka River, near Tbilisi, Georgia (see also [10]).

Age: The fossiliferous organic-rich shales of the Dabakhan Formation date back to the latest part of the Lutetian (e.g., [11]), with an estimated age comprised between 42 and 41 Ma (e.g., [12]).

### 3. *Vinciguerria merklini*

Constrained phylogenetic position: Sister to *Vinciguerria attenuata*

Fossil: *Vinciguerria merklini*, PIN 278/21. The Miocene species *Vinciguerria merklini* from the Tarkhanian Horizon of the Kerch Pensinsula, Crimea is considered to be closely related to the extant *Vinciguerria attenuata* (see [13]).

Age: According to Prokofiev [13], all the material referred to *Vinciguerria merklini* comes from Middle Miocene deposits referred to the Tarkhanian stage of the Eastern Paratethys stratigraphy. Palcu et al. [14] associated this stage to a short flooding event that took place in the Eastern Paratethys between 14.85 and 14.75 Ma.

**4. *Scopeloides violator***

Constrained phylogenetic position: Total group Gonostomatidae (*Margrethia valentinae* + *Cyclothone atraria*)

Fossil: *Scopeloides violator*, MUSE-PAL 2079, from finely laminated black marls of the Chiusole Formation, at Solteri and Monte Solane, Italy. This species, together with its congenerics *Scopeloides bellator* and *Scopeloides* cf. *glarisianus*, documents the existence of the modern gonostomatid body plan in the Early Eocene [15].

Age: Based on its planktonic foraminiferan and calcareous nannoplankton content, the Solteri and Monte Solane sections of the Chiusole Formation have been referred to the Ypresian. The fossiliferous layers of Solteri and Monte Solane are coeval, pertaining to the uppermost portions of the (calcareous nannoplankton) zones NP13 and CNE5 (see [16]), ranging from 49.1 to 48.96 Ma.

**5. *Scopeloides bellator***

Constrained phylogenetic position: Total group Gonostomatidae (*Margrethia valentinae* + *Cyclothone atraria*)

Fossil: *Scopeloides bellator*, MUSE-PAL 12a, from finely laminated black marls of the Chiusole Formation, at Solteri and Monte Solane, Italy. This species, together with its congenerics *Scopeloides violator* and *Scopeloides* cf. *glarisianus*, documents the existence of the modern gonostomatid body plan in the Early Eocene [15,17].

Age: See *Scopeloides violator*.

**6. *Scopeloides* cf. *glarisianus***

Constrained phylogenetic position: Total group Gonostomatidae (*Margrethia valentinae* + *Cyclothone atraria*)

Fossil: *Scopeloides cf. glarisianus*, IGVR 64086–64087, from finely laminated black marls of the Chiusole Formation, at Monte Solane, Italy. This species, together with its congenerics *Scopeloides violator* and *Scopeloides bellator*, documents the existence of the modern gonostomatid body plan in the Early Eocene [15,17].

Age: See *Scopeloides violator*.

**7. *Cyclothone* sp., IUV IL/S7**

Constrained phylogenetic position: Stem *Cyclothone* (*Cyclothone pallida* + *Cyclothone atraria*)

Fossil: *Cyclothone* sp., IUV IL/S7. A still unpublished specimen clearly belonging to the genus *Cyclothone* was collected from the Bartonian laminated marly limestone of the Pabdeh Formation, in the Baba-Heydar locality, Zagros Basin, Iran.

Age: The fish bearing layers of the Pabdeh Formation in the Baba-Heydar locality can be referred to the early Bartonian [18]. Garassino et al. [19] reported the presence of the planktonic foraminiferans *Hantkenina alabamaensis*, *H. compressa* and *Morozovelloides bandyi* in the fish-bearing layers, thereby implying a minimum age comprised between 40.5 and 40.0 Ma [18].

**8. *Cyclothone mukhachevae***

Constrained phylogenetic position: Sister to *Cyclothone atraria*.

Fossil: *Cyclothone mukhachevae*, ZIN 55629. Nazarkin [20] described the fossil bristlemouth species *Cyclothone mukhachevae* from the Miocene Kurasi Formation, Sakhalin Island, Russia and suggested a putative relationship with the extant species *Cyclothone atraria*.

Age: The age of the Kurasi Formation has been established based on diatoms and dinoflagellates that are indicative of the *Denticulopsis simonsenii* Subzone of the *Denticulopsis hyalina* Zone [21,22], corresponding to the late Langhian-early Serravallian, with an estimated age interval of 14.6-13.1 Ma (see [23]).

**9. *Cyclothone gaudanti***

Constrained phylogenetic position: Sister to *Cyclothone signata*.

Fossil: *Cyclothone gaudanti*, NHMW 1999z0042/0020. About 16 articulated skeletons from the laminated marls of the Upper Miocene Makrilia Formation, Ierapetra, Crete, document the existence of the fossil species *Cyclothone gaudanti*, a putative close relative of the extant species *Cyclotone alba*, *Cyclothone braueri* and *Cyclothone signata* [24].

Age: The Tortonian age of theMakrilia Formation has been established based on planktonic foraminiferans and calcareous nannoplankton; the fossiliferous deposits have been referred to NN11a calcareous nannoplankton zone (e.g., [25]), with an age ranging approximately from 8.2 and 7.4 Ma.

**10. *Maurolicus morgani***

Constrained phylogenetic position: Stem *Maurolicus* (*Maurolicus japonicus* + *Maurolicus muelleri*)

Fossil: *Maurolicus morgani*, MNHN-F EiP167. This species described by Arambourg [26] from the laminated marly limestone of the Pabdeh Formation, in the Ilam locality, Zagros Basin, Iran is used to provide a minimum age for the total group *Maurolicus*.

Age: Haghipour & Brants [27] proposed a Middle to Late Eocene age for the fish bearing layers of the Pabdeh Formation exposed in the Ilam locality, western Iran. While a detailed biostratigraphic analysis of these fossiliferous strata is not available, the occurrence of planktonic foraminiferans of the genus *Hantkenina* implies an Eocene age (see [28]). Therefore, we suggest a minimum age of 33.9 Ma (maximum age corresponding to the base of the Priabonian at 37.71 Ma).

**11. *Argyropelecus zagrosensis***

Constrained phylogenetic position: *Argyropelecus affinis* complex (*Argyropelecus affinis* + *Argyropelecus gigas*)

Fossil: *Argyopelecus zagrosensis*, IUV IL/S2. A recently discovered species from the Bartonian laminated marly limestone of the Pabdeh Formation, in the Baba-Heydar locality, Zagros Basin, Iran, is used herein to provide a minimum age for the *Argyropelecus affinis* complex [18].

Age: The fish bearing layers of the Pabdeh Formation in the Baba-Heydar locality can be referred to the early Bartonian [18]. Garassino et al. [19] reported the presence of the planktonic foraminiferans *Hantkenina alabamaensis*, *H. compressa* and *Morozovelloides* *bandyi* in the fish-bearing layers, thereby implying a minimum age comprised between 40.5 and 40.0 Ma [18].

**12. *Argyropelecus iranicus***

Constrained phylogenetic position: *Argyropelecus lychnus* complex (*Argyropelecus aculeatus* + *Argyropelecus sladeni*)

Fossil: *Argyropelecus iranicus* IUV IL/S4. A recently discovered species from the Bartonian laminated marly limestone of the Pabdeh Formation, in the Baba-Heydar locality, Zagros Basin, Iran, is used herein to provide a minimum age for the *Argyropelecus lychnus* complex [18].

Age: The fish bearing layers of the Pabdeh Formation in the Baba-Heydar locality can be referred to the early Bartonian [18]. Garassino et al. [19] reported the presence of the planktonic foraminiferans *Hantkenina alabamaensis*, *H. compressa* and *Morozovelloides bandyi* in the fish-bearing layers, thereby implying a minimum age comprised between 40.5 and 40.0 Ma [18].

**13. *Argyropelecus logearti***

Constrained phylogenetic position: Sister to *Argyropelecus hemigymnus*.

Fossil: *Argyropelecus logearti*, MGPA TOR001. The deep-sea hatchefish *Argyropelecus logearti* is known from the laminated wackestones of the Tufillo Unit (Molisan Units) cropping out at Torricella Peligna, central Italy, as well as from the lower Messinian deposits of northern Italy and Algeria [29,30]. *Argyropelecus logearti* is a Miocene species that forms a sister pair with the extant *Argyropelecus hemigymnus*; this putative sister group relationship is supported by two synapomorphies, presence of a single postabdominal spine and narrow and cyclindrical body trunk [31].

Age: The age of the fish-bearing wackestones of Torricella Peligna has been estimated biostratigraphically, based on planktonic foraminiferans and calcareous nannoplankton [30]. These fossiliferous sediments date back to the Serravallian *Dentoglobigerina altispira altispira* planktonic foraminiferal zone, comprised between about 13.5 and 12.8 Ma [32]. The minimum age of the Messinian material of *Argyropelecus logearti* is about 6.5 Ma.

**14. *Polymetme* sp., (MFM) IY-71**

Constrained phylogenetic position: Stem *Polymetme* (*Polymetme thaeocoryla* + *Polymetme illustris*)

Fossil: *Polymetme* sp., (MFM) IY-71. Well-preserved articulated skeletal remains from the finely laminated tuffaceous sandstones of the Yamami Formation (Morozaki Group) at the Iwaya locality, Aichi Prefecture, Chita Peninsula, Japan, document the existence of the genus *Polymetme* in the Early Miocene [33,34].

Age: The deep-sea fossil-bearing layers of the Yamami Formation have been constrained between 17.4 and 16.6 Ma (Early Miocene, Burdigalian) due to their position around the C5Dn/C5Dr chronozone boundary [35].

**15. *Woodsia* sp., (MFM) MY-3-15 (D-11,12)**

Constrained phylogenetic position: Sister to *Woodsia meyerwardeni*

Fossil: *Woodsia* sp., (MFM) MY-3-15 (D-11,12). Fossil skeletal remains referred to an indeterminate species of the genus *Woodsia* from the laminated tuffaceous sandstones of the Yamami Formation (Morozaki Group), Iwaya locality, Aichi Prefecture, Chita Peninsula, Japan [33,34], are used herein to provide a minimum age for the pair formed by *Woodsia* plus *Phosichthys*.

Age: The deep-sea fossil-bearing layers of the Yamami Formation have been constrained between 17.4 Ma and 16.6 Ma (Early Miocene, Burdigalian) due to their position near the C5Dn/C5Dr chronozone boundary [35].

**16. *Chauliodus* cf. *macoumi***

Constrained phylogenetic position: Sister to *Chauliodus macoumi*.

Fossil: *Chauliodus* cf. *macoumi*, (MFM) MY-Z-3. Articulated skeletal remains referred to *Chauliodus* cf. *macoumi* from the laminated tuffaceous sandstones of the Yamami Formation (Morozaki Group), Iwaya locality, Aichi Prefecture, Chita Peninsula, Japan [33,34], are used herein to document the extinct diversity of the genus *Chauliodus*.

Age: The deep-sea fossil-bearing layers of the Yamami Formation have been constrained between 17.4 Ma and 16.6 Ma (Early Miocene, Burdigalian) due to their position near the C5Dn/C5Dr chronozone boundary [35].

**17. *Chauliodus testa***

Constrained phylogenetic position: Sister to *Chauliodus macoumi* + *Chauliodus* cf. *macoumi*.

Fossil: *Chauliodus testa*, ZIN 55476. Skeletal remains of the extinct species *Chauliodus testa* from the siltstones of the Kurasi Formation exposed along the coastal cliffs of the Tartar Strait, Tomari District, southwestern Sakhalin, Russia [36], are used herein to document the extinct diversity of the genus *Chauliodus*. According to Nazarkin [36], *Chauliodus testa* is extremely similar to *Chauliodus macoumi* from which it differs by having less numerous and completely ossified cervical vertebrae and a reduced predorsal distance.

Age: The age of the Kurasi Formation has been established based on diatoms and dinoflagellates that are indicative of the *Denticulopsis simonsenii* Subzone of the *Denticulopsis hyalina* Zone [21,22], corresponding to the late Langhian-early Serravallian, with an estimated age interval of 14.6-13.1 Ma [23].

**18. *Chauliodus eximius***

Constrained phylogenetic position: *Chauliodus* (*Chauliodus macoumi* + *Chauliodus sloani*)

Fossil: *Chauliodus eximius,* LACM 5244. Crane [37] referred about 44 articulated specimens from the Miocene deposits of the Modelo Formation exposed in the Los Angeles County and Ventura County, California to the extinct species *Chauliodus eximius* that he considered to as closely related to the extant Pacific species *Chauliodus barbatus*.

Age: According to Barron [38], the base and top of the Upper Miocene Modelo Formation range from about 13 to 5.5 Ma.

**19. *Stomias* cf. *affinis***

Constrained phylogenetic position: Sister to *Stomias affinis*

Fossil: *Stomias* cf. *affinis*, LACM 12697. Fink & Fink [39] tentatively referred to *Stomias affinis* a single specimen from the Upper Miocene deposits of the Modelo Formation, Sepulveda Canyon, Los Angeles County, California.

Age: According to Barron [38], the base and top of the Modelo Formation range from about 13 to 5.5 Ma.

**20. *Abruzzoichthys erminioi***

Constrained phylogenetic position: Stem *Astronesthes* (*Astronesthes nigroides* + *Astronesthes illuminatus*)

Fossil: *Abruzzoichthys erminioi*, MGPA TOR 002, from the laminated wackestones of the Tufillo Unit (Molisan Units) cropping out at Torricella Peligna, central Italy. *Abruzzoichthys* is a Miocene genus closely related to *Astronesthes* from which differs by having a discoid posttemporal, greatly elongate body and peculiar relative position of dorsal and anal fins [29]. It is used herein to provide a minimum age of the total group *Astronesthes*.

Age: The age of the fish-bearing wackestones of Torricella Peligna has been estimated biostratigraphically, based on planktonic foraminiferans and calcareous nannoplankton [30]. These fossiliferous sediments date back to the Serravallian *Dentoglobigerina altispira altispira* planktonic foraminiferal zone, comprised between about 13.5 and 12.8 Ma [32].

**Institutional abbreviations:**

IGVR: General inventory of the Verona province (Museo di Storia Naturale di Verona and Antiquarium of San Giorgio di Valpolicella, Verona, Italy); IUV, Museum of Geology, University of Isfahan, Isfahan, Iran; LACM, Natural History Museum of Los Angeles County; MFM, Mizunami Fossil Museum, Mizunami, Japan (the MFM is the repository for numerous fossils from the Yamami Formation collected by the Tokai Fossil Society); MGPA, Museo Geo-Paleontologico dell'Alto Aventino, Palena, Italy; MNHN, Muséum National d'Histoire Naturelle, Paris, France; MUSE, Museo delle Scienze di Trento, Trento, Italy; NHMW, theGeologisch-Paläontologische Abteilung of the Naturhistorisches Museum Wien, Austria; PIN, Borissiak Paleontological Institute, Russian Academy of Sciences, Moscow, ZIN, Zoological Institute, Russian Academy of Sciences, St. Petersburg, Russia.

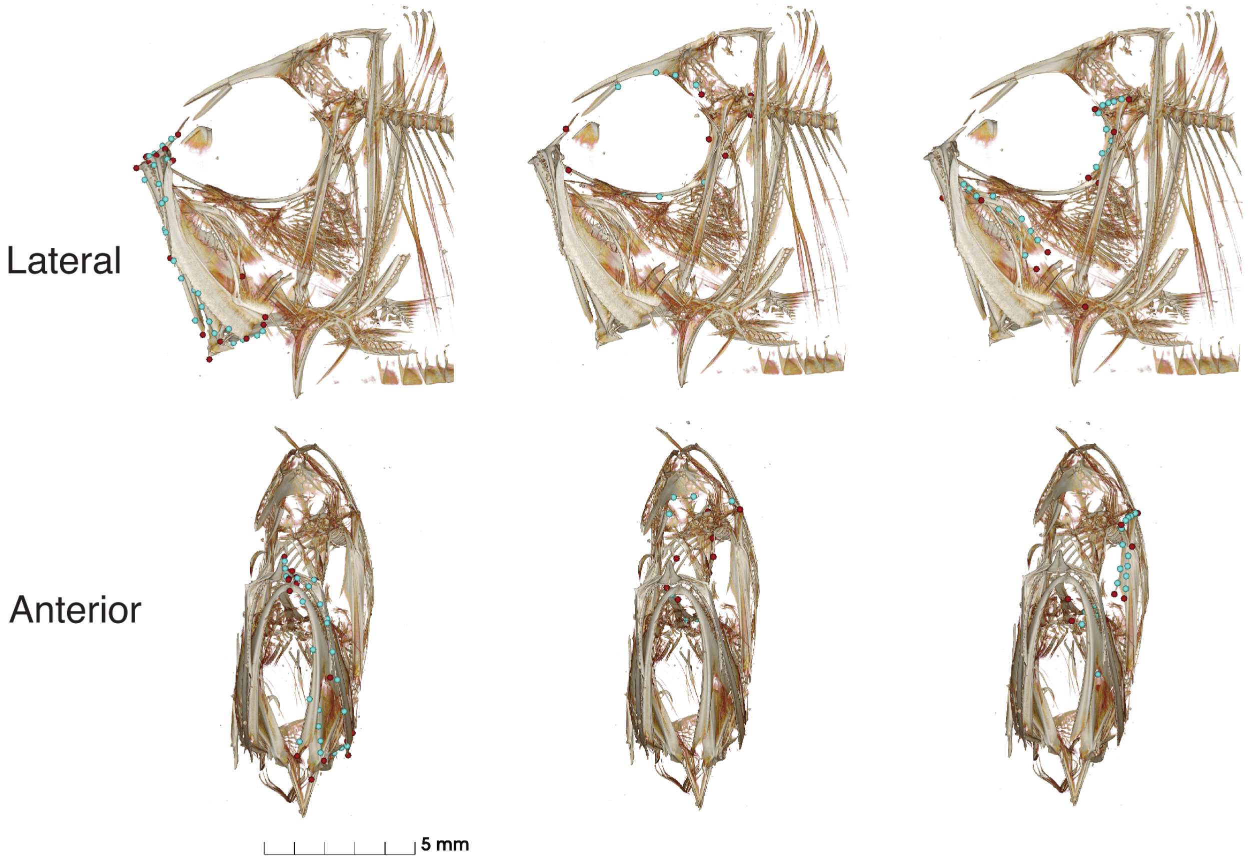


**Figure S1.** Landmarks placed on oral jaws (left), neurocranium (middle), and other bones in skull (right; hyomandibula + hyoid complex + palatine). Red points are fixed landmarks; blue points are semi-sliding landmarks. Scan shown from *Argyropelecus hemigymnus* (UF 180084).

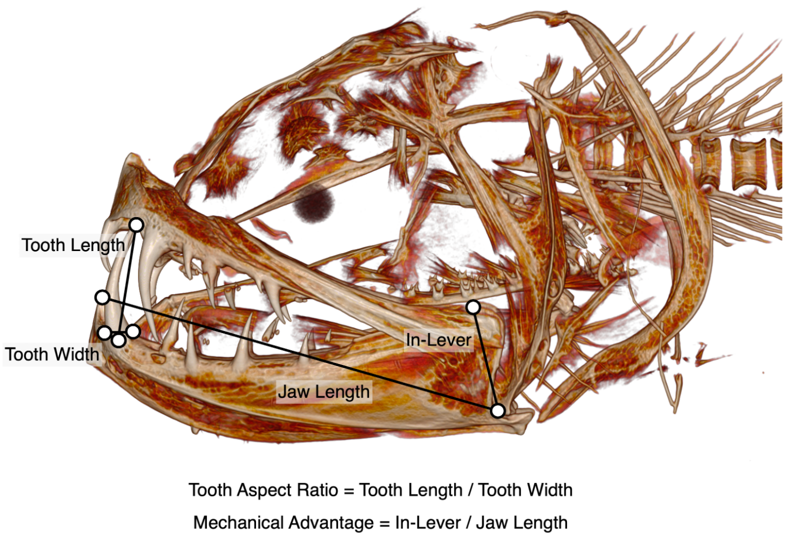


**Figure S2.** Linear measurements of tooth morphology and functional proxies made from CT scans. Scan shown from *Opostomias mitsuii* (LACM 33713.1).

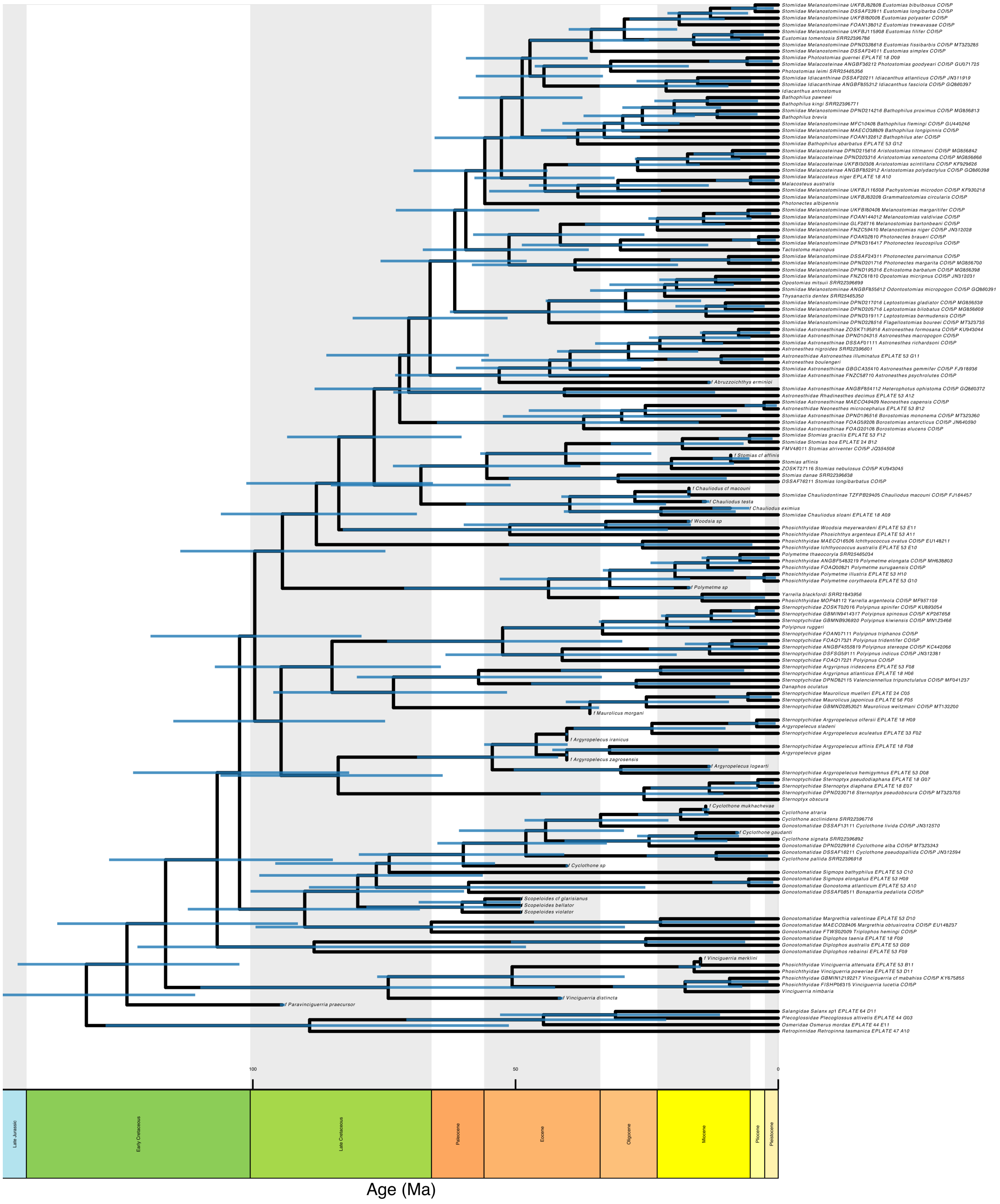


**Figure S3.** Maximum clade credibility tree estimated using FBD showing systematic placement of extinct taxa and 95% highest posterior density around node ages.

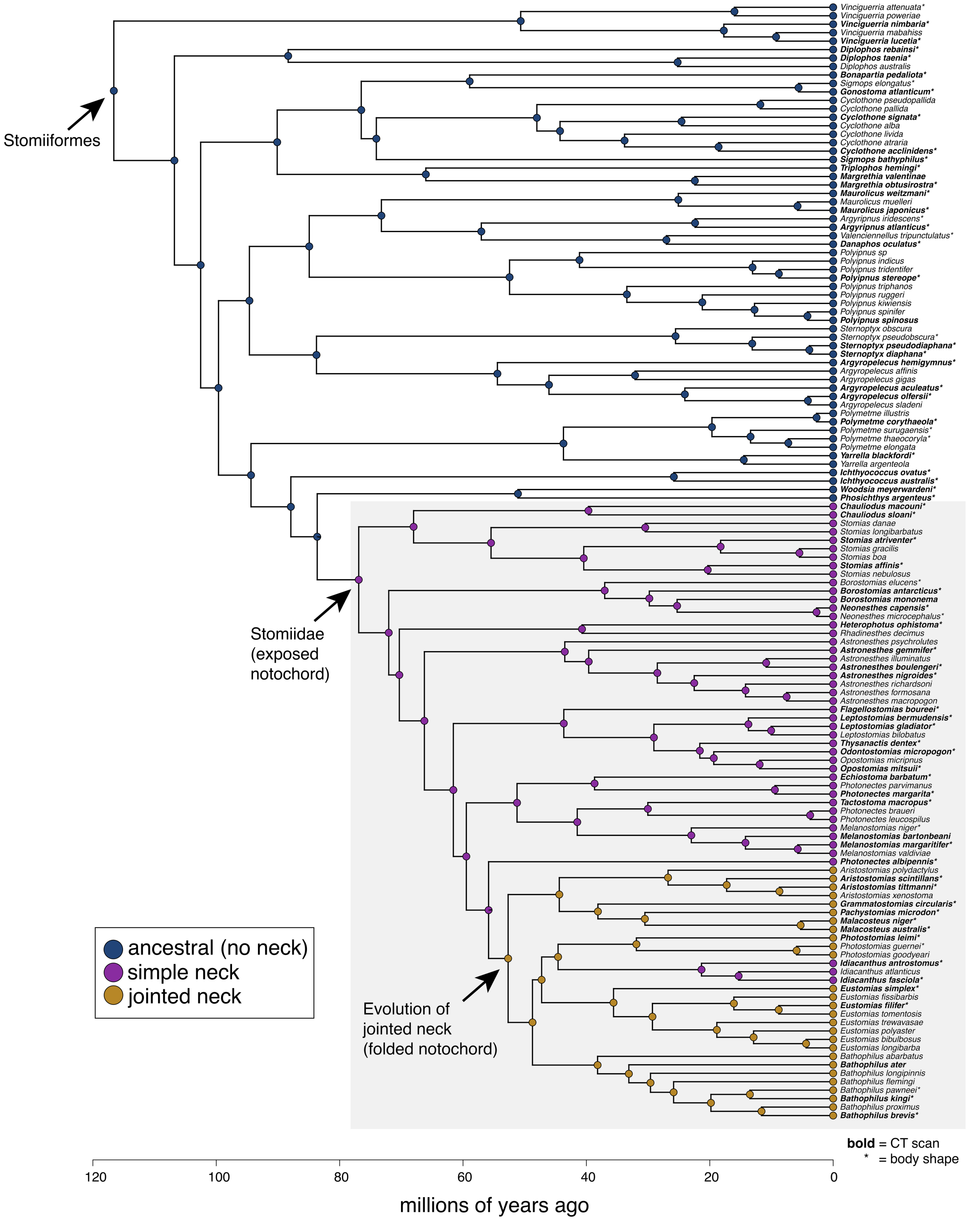


**Figure S4.** SIMMAP ancestral state estimation of neck innovation states, where ancestral = no neck joint, simple = exposed notochord, and jointed = folded notochord. Font style of tip labels indicates overlap with the CT scan and body shape datasets.

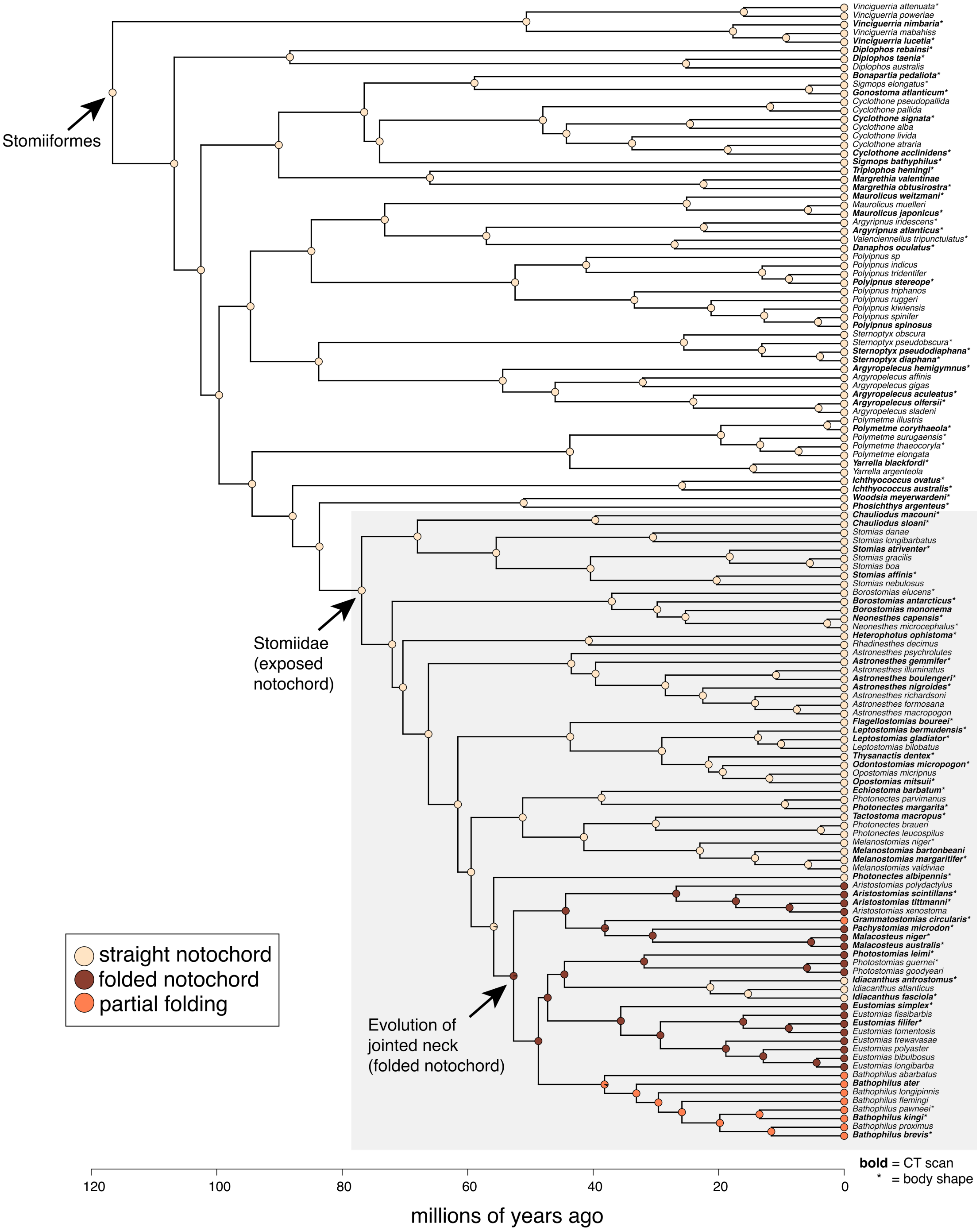


**Figure S5.** SIMMAP ancestral state estimation of the degree of folding of the notochord among taxa in the jointed neck state (Fig. S4). Character states follow Schnell and Johnson (2017). Font style of tip labels indicates overlap with the CT scan and body shape datasets.

**Table S1.** Metadata for the 78 micro-CT scans used in this study, with neck innovation categories.

| Family | Species | Catalog Number | Source | Exposure time | Voltage | Amperage | Media number | Voxel size | Found in phylogeny? | Neck group |
|---|---|---|---|---|---|---|---|---|---|---|
| Gonostomatidae | *Bonapartia pedaliota* | LACM 57237.002 | new scan | 570 | 70 | 214 | 000630578 | 30 um | Y | Ancestral state |
| Gonostomatidae | *Cyclothone acclinidens* | LACM 44870.005 | new scan | 570 | 53 | 165 | 000630294 | 0.024 mm | Y | Ancestral state |
| Gonostomatidae | *Cyclothone microdon* | SIO 75-450 | new scan | 420 | 60 | 160 | 000723587 | 20 um | N | Ancestral state |
| Gonostomatidae | *Cyclothone signata* | LACM 30509.030 | new scan | 470 | 55 | 200 | 000678868 | 0.017 mm | Y | Ancestral state |
| Gonostomatidae | *Diplophos rebainsi* | SIO 61-42 | new scan | 420 | 58 | 165 | 000723859 | 48 um | Y | Ancestral state |
| Gonostomatidae | *Diplophos taenia* | SIO 00-177 | new scan | 420 | 50 | 160 | 000723178 | 30 um | Y | Ancestral state |
| Gonostomatidae | *Gonostoma atlanticum* | LACM 36301.002 | new scan | 570 | 40 | 165 | 000664075 | 26.5 um | Y | Ancestral state |
| Gonostomatidae | *Gonostoma bathyphilum* | LACM 11546.010 | new scan | 470 | 60 | 200 | 000678455 | 0.027 mm | *Sigmops bathyphilus* | Ancestral state |
| Gonostomatidae | *Margrethia obtusirostra* | SIO 77-167 | new scan | 420 | 60 | 140 | 000716089 | 23 um | Y | Ancestral state |
| Gonostomatidae | *Margrethia valentinae* | LACM 11274.21 | new scan | 570 | 60 | 210 | 000631443 | 0.024 mm | Y | Ancestral state |
| Gonostomatidae | *Triplophos hemingi* | UF 237090 | new scan | 420 | 60 | 200 | 000710488 | 27.5 um | Y | Ancestral state |
| Phosichthyidae | *Ichthyococcus irregularis* | LACM 55959.007 | new scan | 470 | 60 | 200 | 000637776 | 0.026 mm | *Ichthyococcus australis* | Ancestral state |
| Phosichthyidae | *Ichthyococcus ovatus* | SIO 76-81 | new scan | 420 | 60 | 180 | 000723502 | 20 um | Y | Ancestral state |
| Phosichthyidae | *Phosichthys sp.* | LACM 11505.19 | new scan | 570 | 40 | 214 | 000632350 | 0.03 mm | *Phosichthys argenteus* | Ancestral state |
| Phosichthyidae | *Pollichthys mauli* | SIO 70-334 | new scan | 420 | 57 | 160 | 000721878 | 15 um | N | Ancestral state |
| Phosichthyidae | *Polymetme corythaeola* | LACM 44730 | new scan | 570 | 70 | 214 | 000631403 | 0.03 mm | Y | Ancestral state |
| Phosichthyidae | *Vinciguerria lucetia* | LACM 30040.029 | new scan | 570 | 40 | 165 | 000630235 | 0.03 mm | Y | Ancestral state |
| Phosichthyidae | *Vinciguerria nimbaria* | LACM 57315.022 | new scan | 470 | 61 | 200 | 000678504 | 0.020 mm | Y | Ancestral state |
| Phosichthyidae | *Woodsia sp.* | LACM 11542.010 | new scan | 570 | 40 | 165 | 000630300 | 0.03 mm | *Woodsia meyerwardeni* | Ancestral state |
| Phosichthyidae | *Yarrella blackfordi* | LACM 36174.001 | new scan | 570 | 55 | 214 | 000633313 | 0.03 mm | Y | Ancestral state |
| Sternoptychidae | *Argyripnus ephippiatus* | SIO 00-20 | Morphosource | 0.200098 | 60 | 200 | 000073428 | 0.018195 mm | *Argyripnus atlanticus* | Ancestral state |
| Sternoptychidae | *Argyropelecus aculeatus* | SIO 70-314 | new scan | 420 | 60 | 210 | 000718912 | 23 um | Y | Ancestral state |

| | | | | | | | | | | | |
|---|---|---|---|---|---|---|---|---|---|---|---|
| Sternoptychidae | *Argyropelecus hemigymnus* | UF 180084 | Morphosource | 0.250097 | 80 | 200 | 000437872 | 0.009954 mm | Y | | Ancestral state |
| Sternoptychidae | *Argyropelecus lychnus* | SIO 76-54 | new scan | not given | 60 | 220 | 000723285 | 29 um | N | | Ancestral state |
| Sternoptychidae | *Argyropelecus olfersii* | LACM 10199 | new scan | 500 | 90 | 159 | 000628915 | 22 um | Y | | Ancestral state |
| Sternoptychidae | *Danaphos oculatus* | LACM 30611.10 | new scan | 570 | 53 | 170 | 000633288 | 0.024 mm | Y | | Ancestral state |
| Sternoptychidae | *Maurolicus amethystinopunctatus* | MCZ 152925 | Morphosource | 1.35 | 40 | 200 | 000088389 | 0.023072 mm | N | | Ancestral state |
| Sternoptychidae | *Maurolicus australis* | LACM 10989 | new scan | 470 | 55 | 180 | 000678426 | 0.018 mm | N | | Ancestral state |
| Sternoptychidae | *Maurolicus japonicus* | MCZ 31487 | new scan | 410 | 60 | 160 | 000693215 | 0.027 mm | Y | | Ancestral state |
| Sternoptychidae | *Maurolicus parvipinnis* | LACM 10365.004 | new scan | 570 | 40 | 160 | 000630585 | 25 um | N | | Ancestral state |
| Sternoptychidae | *Maurolicus wietzmani* | LACM 61404 | new scan | 570 | 53 | 165 | 000691373 | 20 um | Y | | Ancestral state |
| Sternoptychidae | *Polyipnus clarus* | UF 178351 | new scan | 470 | 60 | 180 | 000710368 | 25 um | *Polyipnus spinosus* | | Ancestral state |
| Sternoptychidae | *Polyipnus stereope* | LACM 36062.004 | new scan | 470 | 60 | 200 | 000678882 | 0.020 mm | Y | | Ancestral state |
| Sternoptychidae | *Sternoptyx diaphana* | LACM 61775 | new scan | 570 | 53 | 165 | 000631421 | 0.021 mm | Y | | Ancestral state |
| Sternoptychidae | *Sternoptyx pseudodiaphana* | SIO 84-13 | new scan | 420 | 60 | 185 | 000723594 | 20 um | Y | | Ancestral state |
| Stomiidae | *Aristostomias scintillans* | LACM 9077.3 | new scan | 470 | 62 | 200 | 000634645 | 0.03 mm | Y | | Jointed neck |
| Stomiidae | *Aristostomias tittmanni* | MCZ 70515-b | new scan | 450 | 60 | 160 | 000693597 | 0.03 mm | Y | | Jointed neck |
| Stomiidae | *Astronesthes boulengeri* | LACM 10273.000 | new scan | 470 | 60 | 200 | 000678889 | 0.030 mm | Y | | Simple neck |
| Stomiidae | *Astronesthes gemmifer* | SIO 71-71 | new scan | 420 | 58 | 160 | 000723267 | 37 um | Y | | Simple neck |
| Stomiidae | *Astronesthes nigroides* | SIO 08-45 | new scan | 420 | 60 | 165 | 000723441 | 25 um | Y | | Simple neck |
| Stomiidae | *Astronesthes trifibulatus* | LACM 56419.5 | new scan | 470 | 50 | 200 | 000634651 | 0.026 mm | N | | Simple neck |
| Stomiidae | *Bathophilus ater* | LACM 11231 | new scan | 570 | 50 | 200 | 000633308 | 0.03 mm | Y | | Jointed neck |
| Stomiidae | *Bathophilus brevis* | LACM 44154.001 | new scan | 470 | 65 | 200 | 000678875 | 0.025 mm | Y | | Jointed neck |
| Stomiidae | *Bathophilus kingi* | SIO 97-92 | new scan | 420 | 60 | 170 | 000723260 | 25 um | Y | | Jointed neck |
| Stomiidae | *Bathophilus melas* | MCZ 84884 | Morphosource | 0.35 | 40 | 200 | 000066574 | 0.056840 mm | N | | Jointed neck |
| Stomiidae | *Borostomias antarcticus* | SIO 75-451 | new scan | 420 | 60 | 200 | 000723217 | 30 um | Y | | Simple neck |
| Stomiidae | *Borostomias mononema* | UF 172459 | Morphosource | 0.200098 | 60 | 220 | 000051431 | 0.070717 mm | Y | | Simple neck |
| Stomiidae | *Chauliodus barbatus* | SIO 68-536 | new scan | 420 | 55 | 150 | 000723922 | 32 um | N | | Simple neck |

| | | | | | | | | | | |
|---|---|---|---|---|---|---|---|---|---|---|
| Stomiidae | *Chauliodus macouni* | LACM 8989.030 | new scan | 470 | 60 | 200 | 000636366 | 0.03 mm | Y | Simple neck |
| Stomiidae | *Chauliodus sloani* | SIO 84-53 | new scan | 420 | 58 | 165 | 000723792 | 38 um | Y | Simple neck |
| Stomiidae | *Echiostoma barbatum* | LACM 56419.22 | new scan | 470 | 54 | 200 | 000634669 | 0.026 mm | Y | Simple neck |
| Stomiidae | *Eustomias filifer* | MCZ 153128 | new scan | 410 | 55 | 160 | 000693202 | 0.026 mm | Y | Jointed neck |
| Stomiidae | *Eustomias jimcraddocki* | MCZ 163108 | new scan | 450 | 60 | 160 | 000693756 | 0.027 mm | *Eustomias simplex* | Jointed neck |
| Stomiidae | *Eustomias schiffi* | MCZ 169565 | new scan | 450 | 55 | 160 | 000693248 | 0.024 mm | N | Jointed neck |
| Stomiidae | *Flagellostomias boureei* | LACM 57242.002 | new scan | 470 | 60 | 200 | 000636903 | 0.03 mm | Y | Simple neck |
| Stomiidae | *Grammatostomias circularis* | MCZ 163247 | new scan | 410 | 55 | 160 | 000693222 | 0.022 mm | Y | Jointed neck |
| Stomiidae | *Heterophotus ophistoma* | LACM 54114.025 | new scan | 570 | 60 | 200 | 000633815 | 0.03 mm | Y | Simple neck |
| Stomiidae | *Idiacanthus antrostomus* | LACM 30263.007 | new scan | 470 | 50 | 200 | 000635308 | 0.022 mm | Y | Simple neck |
| Stomiidae | *Idiacanthus fasciola* | SIO 70-341 | new scan | 317 | 48 | 170 | 000724958 | 30 um | Y | Simple neck |
| Stomiidae | *Leptostomias gladiator* | SIO 97-39 | new scan | 420 | 48 | 160 | 000724939 | 31 um | Y | Simple neck |
| Stomiidae | *Leptostomias macronema* | SIO 75-634 | new scan | 420 | 47 | 150 | 000724587 | 28 um | *Leptostomias bermudensis* | Simple neck |
| Stomiidae | *Malacosteus australis* | SIO 61-36 | new scan | 420 | 52 | 180 | 000724611 | 30 um | Y | Jointed neck |
| Stomiidae | *Malacosteus niger* | LACM 57120.006 | new scan | 470 | 65 | 200 | 000636726 | 0.024 mm | Y | Jointed neck |
| Stomiidae | *Melanostomias bartonbeani* | SIO 61-36 | new scan | 420 | 56 | 180 | 000724975 | 42 um | Y | Simple neck |
| Stomiidae | *Melanostomias cf. margaritifer* | SIO 70-314 | new scan | 420 | 56 | 160 | 000722663 | 38 um | *Melanostomias margaritifer* | Simple neck |
| Stomiidae | *Melanostomias tentaculatus* | LACM 10362.006 | new scan | 470 | 60 | 200 | 000636336 | 0.03 mm | N | Simple neck |
| Stomiidae | *Neonesthes capensis* | YPM 025806 | Morphosource | 0.200098 | 70 | 200 | 000426441 | 0.030597 mm | Y | Simple neck |
| Stomiidae | *Odontostomias micropogon* | SIO 71-382 | new scan | 420 | 50 | 150 | 000724164 | 30 um | Y | Simple neck |
| Stomiidae | *Opostomias mitsuii* | LACM 33713.1 | new scan | 570 | 60 | 214 | 000632361 | 0.03 mm | Y | Simple neck |
| Stomiidae | *Pachystomias microdon* | LACM 54114.04 | new scan | 470 | 65 | 200 | 000636731 | 0.03 mm | Y | Jointed neck |
| Stomiidae | *Photonectes albipennis* | YPM 010032 | Morphosource | 0.200098 | 70 | 200 | 000426701 | 0.017716 mm | Y | Simple neck |
| Stomiidae | *Photonectes margarita* | LACM 9576.023 | new scan | 470 | 60 | 200 | 000635291 | 0.026 mm | Y | Simple neck |
| Stomiidae | *Photostomias liemi* | LACM 36130.008 | new scan | 470 | 60 | 200 | 000636909 | 0.03 mm | Y | Jointed neck |

| | | | | | | | | | | |
|---|---|---|---|---|---|---|---|---|---|---|
| Stomiidae | *Stomias affinis* | SIO 69-23 | new scan | 420 | 55 | 150 | 000723277 | 37 um | Y | Simple neck |
| Stomiidae | *Stomias atriventer* | LACM 30056.005 | new scan | 420 | 73 | 115 | 000727081 | 32 um | Y | Simple neck |
| Stomiidae | *Tactostoma macropus* | LACM 9716.004 | new scan | 470 | 50 | 200 | 000634764 | 0.03 mm | Y | Simple neck |
| Stomiidae | *Thysanactis dentex* | YPM 025683 | Morphosource | 0.200098 | 70 | 200 | 000426421 | 0.112868 mm | Y | Simple neck |
| Stomiidae | *Trigonolampa miriceps* | YPM 011438 | Morphosource | 0.200098 | 80 | 200 | 000426709 | 0.042691 mm | N | Simple neck |

**Table S2.** Description of 102 landmarks used for this study and comparison with the scheme used in Miller et al. [40].

| **Landmark number** | **Bone** | **Landmark type** | **Description** | **Usage in Miller et al. [40]?** |
|---|---|---|---|---|
| 1 | Premaxilla | fixed | Tip of most medial tooth on premaxilla | same |
| 2 | Premaxilla | fixed | Base of most medial tooth on premaxilla | same |
| 3 | Premaxilla | fixed | Anteriormost medial point on dentigerous arm of premaxilla | modified placement |
| 4 | Premaxilla | fixed | Proximal vertex between ascending and descending processes of premaxilla | same |
| 5 | premaxilla | fixed | Most distal point of the premaxilla | same |
| 6 | dentary | fixed | Tip of the most medial tooth on dentary | same |
| 7 | dentary | fixed | Posterior-dorsalmost tip of dentary | same |
| 8 | dentary | fixed | Ventral tip of medial dentary symphysis | same |
| 9 | anguloarticular | fixed | Anterior-distal tip of the anguloarticular | same |
| 10 | anguloarticular | fixed | Tip of ascending process of anguloarticular | same |
| 11 | dentary | fixed | Ventral-posterior point of dentary | same |
| 12 | anguloarticular | fixed | Center of jaw joint on the quadrate | same |
| 13 | anguloarticular | fixed | Posterior tip of retroarticular (or articular if retroarticular is absent) | same |
| 14 | neurocranium | fixed | Anterior medial point of vomer | same |
| 15 | neurocranium | fixed | Lateral anterior tip of vomer | same |
| 16 | neurocranium | fixed | Proximal lateral ethmoid-parasphenoid margin | same |
| 17 | neurocranium | fixed | Anterior point of orbit (determined functionally, not anatomically) | modified placement |
| 18 | neurocranium | fixed | Posterior point of orbit (determined functionally, not anatomically) | modified placement |
| 19 | neurocranium | fixed | Pterosphenoid-parasphenoid margin | same |
| 20 | neurocranium | fixed | Midpoint of parasphenoid-basioccipital margin | same |
| 21 | neurocranium | fixed | Prootic foramen | same |
| 22 | neurocranium | fixed | Dorsal-most point of articulation between basioccipital and first vertebrate | same |
| 23 | neurocranium | fixed | Ventral-most point of articulation between basioccipital and first vertebrate | same |
| 24 | premaxilla | sliding | Curve along ascending process of premaxilla (if ascending process is missing, placed on medial symphysis of premaxillae) [point 1, nearest landmark #**2**] | same |
| 25 | premaxilla | sliding | Curve along ascending process of premaxilla (if ascending process is missing, placed on medial symphysis of premaxillae) [point 2] | same |
| 26 | premaxilla | sliding | Curve along ascending process of premaxilla (if ascending process is missing, placed on medial symphysis of premaxillae) [point 3, nearest landmark #3] | same |
| 27 | dentary | sliding | Curve along ventral margin of dentary [point 1, nearest landmark #8] | same |
| 28 | dentary | sliding | Curve along ventral margin of dentary [point 2] | same |

| | | | | |
|---|---|---|---|---|
| 29 | dentary | sliding | Curve along ventral margin of dentary [point 3] | same |
| 30 | dentary | sliding | Curve along ventral margin of dentary [point 4, nearest landmark #11] | same |
| 31 | neurocranium | sliding | Curve along ventral margin of parasphenoid [point 1, nearest landmark #14] | same |
| 32 | neurocranium | sliding | Curve along ventral margin of parasphenoid [point 2] | same |
| 33 | neurocranium | sliding | Curve along ventral margin of parasphenoid [point 3] | same |
| 34 | neurocranium | sliding | Curve along ventral margin of parasphenoid [point 4, nearest landmark #20] | same |
| 35 | neurocranium | sliding | Curve along dorsal surface of orbit [point 1, nearest landmark #17] | modified placement |
| 36 | neurocranium | sliding | Curve along dorsal surface of orbit [point 2] | modified placement |
| 37 | neurocranium | sliding | Curve along dorsal surface of orbit [point 3] | modified placement |
| 38 | neurocranium | sliding | Curve along dorsal surface of orbit [point 4, nearest landmark #18] | modified placement |
| 39 | maxilla | fixed | Distal ventral anterior margin of maxilla | same |
| 40 | maxilla | fixed | Distal ventral posterior margin of maxilla | same |
| 41 | maxilla | fixed | Proximal dorsal anterior margin of maxilla | same |
| 42 | maxilla | fixed | Proximal dorsal posterior margin of maxilla | same |
| 43 | hyomandibula | fixed | Anterior dorsal-most point of hyomandibula | same |
| 44 | hyomandibula | fixed | Posterior-most point of contact on the distal face of the hyomandibula between the hyomandibula and the pterotic | same |
| 45 | hyomandibula | fixed | Distal-most point of lateral projection of hyomandibula | same |
| 46 | hyomandibula | fixed | Anterior ventral end of the vertical limb of hyomandibula | same |
| 47 | hyomandibula | fixed | Ventral posterior-most point of hyomandibula | same |
| 48 | anguloarticular | sliding | Curve along ascending process of the anguloarticular [point 1, nearest landmark #10] | same |
| 49 | anguloarticular | sliding | Curve along ascending process of the anguloarticular [point 2] | same |
| 50 | anguloarticular | sliding | Curve along ascending process of the anguloarticular [point 3] | same |
| 51 | anguloarticular | sliding | Curve along ascending process of the anguloarticular [point 4, nearest landmark #12] | same |
| 52 | anguloarticular | sliding | Curve along lateral process of the anguloarticular [point 1, nearest landmark #9] | same |
| 53 | anguloarticular | sliding | Curve along lateral process of the anguloarticular [point 2] | same |
| 54 | anguloarticular | sliding | Curve along lateral process of the anguloarticular [point 3] | same |
| 55 | anguloarticular | sliding | Curve along lateral process of the anguloarticular [point 4, nearest landmark #12] | same |
| 56 | premaxilla | sliding | Curve along lateral process of the premaxilla [point 1, nearest landmark #2] | same |
| 57 | premaxilla | sliding | Curve along lateral process of the premaxilla [point 2] | same |
| 58 | premaxilla | sliding | Curve along lateral process of the premaxilla [point 3] | same |
| 59 | premaxilla | sliding | Curve along lateral process of the premaxilla [point 4] | same |
| 60 | premaxilla | sliding | Curve along lateral process of the premaxilla [point 5, nearest landmark #5] | same |

| | | | | |
|---|---|---|---|---|
| 61 | hyomandibula | sliding | Curve along anterior face of the hyomandibula [point 1, nearest landmark #44] | same |
| 62 | hyomandibula | sliding | Curve along anterior face of the hyomandibula [point 2] | same |
| 63 | hyomandibula | sliding | Curve along anterior face of the hyomandibula [point 3] | same |
| 64 | hyomandibula | sliding | Curve along anterior face of the hyomandibula [point 4, nearest landmark #47] | same |
| 65 | hyomandibula | sliding | Curve along dorsal surface of the hyomandibula [point 1, nearest landmark #44] | same |
| 66 | hyomandibula | sliding | Curve along dorsal surface of the hyomandibula [point 2] | same |
| 67 | hyomandibula | sliding | Curve along dorsal surface of the hyomandibula [point 3] | same |
| 68 | hyomandibula | sliding | Curve along dorsal surface of the hyomandibula [point 4, nearest landmark #45] | same |
| 69 | hyomandibula | sliding | Curve along posterior face of hyomandibula [point 1, nearest landmark #46] | same |
| 70 | hyomandibula | sliding | Curve along posterior face of hyomandibula [point 2] | same |
| 71 | hyomandibula | sliding | Curve along posterior face of hyomandibula [point 3] | same |
| 72 | hyomandibula | sliding | Curve along posterior face of hyomandibula [point 4, nearest landmark #48] | same |
| 73 | hyoid complex | fixed | Anterior ventral-most point of the anterior ceratohyal [ceratohyal] | same |
| 74 | hyoid complex | fixed | Anterior dorsal-most point of the anterior ceratohyal [ceratohyal] | same |
| 75 | hyoid complex | fixed | Dorsal margin of the anterior ceratohyal [ceratohyal] and posterior ceratohyal [epihyal] | same |
| 76 | hyoid complex | fixed | Posterior-most tip of the posterior ceratohyal [epihyal] | same |
| 77 | hyoid complex | fixed | Ventral margin of the anterior ceratohyal [ceratohyal] and posterior ceratohyal [epihyal] | same |
| 78 | hyoid complex | sliding | Curve along dorsal surface of the anterior ceratohyal [point 1, nearest landmark #75] | same |
| 79 | hyoid complex | sliding | Curve along dorsal surface of the anterior ceratohyal [point 2] | same |
| 80 | hyoid complex | sliding | Curve along dorsal surface of the anterior ceratohyal [point 3] | same |
| 81 | hyoid complex | sliding | Curve along dorsal surface of the anterior ceratohyal [point 4] | same |
| 82 | hyoid complex | sliding | Curve along dorsal surface of the anterior ceratohyal [point 5, nearest landmark #76] | same |
| 83 | hyoid complex | sliding | Curve along ventral surface of the anterior ceratohyal [point 1, nearest landmark #74] | same |
| 84 | hyoid complex | sliding | Curve along ventral surface of the anterior ceratohyal [point 2] | same |
| 85 | hyoid complex | sliding | Curve along ventral surface of the anterior ceratohyal [point 3] | same |
| 86 | hyoid complex | sliding | Curve along ventral surface of the anterior ceratohyal [point 4] | same |
| 87 | hyoid complex | sliding | Curve along ventral surface of the anterior ceratohyal [point 5, nearest landmark #78] | Same` |
| 88 | palatine | fixed | Medial anterior-most tip of palatine | not used |
| 89 | palatine | fixed | Medial posterior-most tip of palatine | not used |
| 90 | palatine | sliding | Curve along medial edge of palatine [point 1, nearest landmark #89] | not used |
| 91 | palatine | sliding | Curve along medial edge of palatine [point 2] | not used |

| | | | | |
|---|---|---|---|---|
| 92 | palatine | sliding | Curve along medial edge of palatine [point 3] | not used |
| 93 | palatine | sliding | Curve along medial edge of palatine [point 4] | not used |
| 94 | palatine | sliding | Curve along medial edge of palatine [point 5, nearest landmark #90] | not used |
| 95 | palatine | fixed | Tip of the anterior-most palatine tooth | not used |
| 96 | palatine | fixed | Base of the anterior-most palatine tooth | not used |
| 97 | maxilla | sliding | Curve along ventral surface of the maxilla [point 1, nearest landmark #42] | not used |
| 98 | maxilla | sliding | Curve along ventral surface of the maxilla [point 2] | not used |
| 99 | maxilla | sliding | Curve along ventral surface of the maxilla [point 3] | not used |
| 100 | maxilla | sliding | Curve along ventral surface of the maxilla [point 4] | not used |
| 101 | maxilla | sliding | Curve along ventral surface of the maxilla [point 5, nearest landmark #40] | not used |
| 102 | premaxilla | fixed | Dorsal apex of the structure on the premaxilla that interacts with the maxilla | not used |

**Table S3.** Likelihood ratio test comparing Mk models of transitions among neck and notochord folding categories. Best-fitting models are in bold.

| Analysis | Model | log(L) | Degrees of freedom | AIC | AIC weight |
|---|---|---|---|---|---|
| Neck types (Fig. S4) | **fit_ER** | **-22.04** | **1** | **46.07** | **0.50** |
| | fit_SYM | -20.12 | 3 | 46.25 | 0.46 |
| | fit_ARD | -19.50 | 6 | 51.00 | 0.04 |
| Degree of notochord folding (Fig. S5) | fit_ER | -25.82 | 1 | 53.64 | 0.10 |
| | **fit_SYM** | **-21.87** | **3** | **49.73** | **0.69** |
| | fit_ARD | -20.05 | 6 | 52.10 | 0.21 |

**Table S4 a-b.** Significance of rate differences among the three groups (Jointed neck, Simple neck, Ancestral state) across phenotypes using 'compare.evol.rates' function in geomorph [41]. Significant p-values in bold.

A, Significance of rate differences among groups.

| Phenotype | P value | Effect size |
|---|---|---|
| Whole skull | **0.0001** | 4.02 |
| Oral jaws | 0.1017 | 1.32 |
| Neurocranium | **0.0001** | 3.83 |
| Body | **0.0332** | 1.82 |
| Teeth | **0.0006** | 2.79 |

B, Values of rates and disparity plotted in Fig. 3. See Table S5 for statistical significance of disparity.

| Phenotype | Group | Rate | Disparity |
|---|---|---|---|
| Whole skull | Jointed neck | 0.00000016 | 0.0014 |
| | Simple neck | 0.00000006 | 0.0011 |
| | Ancestral (no neck) | 0.00000007 | 0.0015 |
| Oral jaws | Jointed neck | 0.00000013 | 0.0008 |
| | Simple neck | 0.00000013 | 0.0012 |
| | Ancestral (no neck) | 0.00000016 | 0.0014 |
| Neurocranium | Jointed neck | 0.00002316 | 0.0512 |
| | Simple neck | 0.00000743 | 0.0240 |
| | Ancestral (no neck) | 0.00000834 | 0.0458 |
| Body | Jointed neck | 0.01477581 | 7.9540 |
| | Simple neck | 0.01099598 | 9.1961 |
| | Ancestral (no neck) | 0.00961418 | 14.2322 |
| Teeth | Jointed neck | 0.00450072 | 1.2061 |
| | Simple neck | 0.01400051 | 3.8346 |
| | Ancestral (no neck) | 0.00416077 | 2.4737 |

**Table S5.** Significance in morphological disparity among the three groups across phenotypes using the 'morphol.disparity' function in geomorph [41]. Significant p-values in bold.

Whole skull:

| | Jointed neck | Simple neck | Ancestral state |
|---|---|---|---|
| Jointed neck | NA | | |
| Simple neck | 0.35 | NA | |
| Ancestral state | 0.83 | 0.13 | NA |

Oral jaws:

| | Jointed neck | Simple neck | Ancestral state |
|---|---|---|---|
| Jointed neck | NA | | |
| Simple neck | 0.36 | NA | |
| Ancestral state | 0.14 | 0.60 | NA |

Neurocranium:

| | Jointed neck | Simple neck | Ancestral state |
|---|---|---|---|
| Jointed neck | NA | | |
| Simple neck | **0.01** | NA | |
| Ancestral state | 0.60 | **0.01** | NA |

Body:

| | Jointed neck | Simple neck | Ancestral state |
|---|---|---|---|
| Jointed neck | NA | | |
| Simple neck | 0.66 | NA | |
| Ancestral state | **0.02** | **0.01** | NA |

Teeth:

| | Jointed neck | Simple neck | Ancestral state |
|---|---|---|---|
| Jointed neck | NA | | |
| Simple neck | **0.02** | NA | |
| Ancestral state | 0.26 | 0.13 | NA |

**Table S6.** Innovations found in the neck-associated stomiid genera. Characters from refs. [42,43]. Subfamily classification from ref. [44].

| Genus | Traditional subfamily | Folding of notochord | Intramandibular membrane | Long-wave (red) bioluminescence and vision |
|---|---|---|---|---|
| *Aristostomias* | Malacosteinae | Folded | Absent | Yes |
| *Malacosteus* | Malacosteinae | Folded | Absent | Yes |
| *Photostomias* | Malacosteinae | Folded | Absent | No |
| *Bathophilus* | Melanostomiinae | Partially folded | Present | No |
| *Eustomias* | Melanostomiinae | Folded | Present | No |
| *Grammatostomias* | Melanostomiinae | Partially folded | Present | No |
| *Pachystomias* | Melanostomiinae | Folded | Present | Yes |
